# Cellular, ultrastructural and molecular analyses of epidermal cell development in the planarian *Schmidtea mediterranea*

**DOI:** 10.1101/179986

**Authors:** Li-Chun Cheng, Kimberly C. Tu, Chris W. Seidel, Sofia M.C. Robb, Fengli Guo, Alejandro Sánchez Alvarado

## Abstract

The epidermis is essential for animal survival, providing both a protective barrier and cellular sensor to external environments. The generally conserved embryonic origin of the epidermis, but the broad morphological and functional diversity of this organ across animals is puzzling. We define the transcriptional regulators underlying epidermal lineage differentiation in the planarian *Schmidtea mediterranea,* an invertebrate organism that, unlike fruitflies and nematodes, continuously replaces its epidermal cells. We find that *Smed-p53, Sox* and *Pax* transcription factors are essential regulators of epidermal homeostasis, and act cooperatively to regulate genes associated with early epidermal precursor cell differentiation, including a tandemly arrayed novel gene family *(prog)* of secreted proteins. Additionally, we report on the discovery of distinct and previously undescribed secreted organelles whose production is dependent on the transcriptional activity of *soxP-3,* and which we term Hyman vesicles.

## Introduction

The epidermis is a central innovation in the evolutionary history of animals. This outer cell layer not only serves as the first line of defense to environmental vicissitudes such as injury and infection, but also plays diverse roles including regulation of body temperature and water balance, respiration, sensing the surroundings, and producing signals related to behavior (Tyler, 1984). Because of its complexity and constant environmental interactions, epidermal cellular composition, architecture and homeostasis are remarkably diverse both morphologically and biochemically throughout the animal kingdom (Tyler, 1984). The planarian adult epidermis is a monostratified tissue that consists of a single layer of post-mitotic, differentiated, multi-ciliated and non-ciliated cell types (Rompolas et al., 2010). This non-cuticularized epidermis sits atop a basement membrane overlying the body wall musculature (Hyman, 1951). Several past andrecent studies have shown that the planarian epidermis is composed of a remarkable diversity of differentiated cell types (McGee et al., 1997; Tyler and Tyler, 1997; Liana et al., 2012; Tu et al., 2015). Moreover, these differentiated cells are continuously replaced by post-mitotic epithelial precursors derived from a pool of dividing pluripotent stem cells known as neoblasts that reside deep in the mesenchyme (Newmark and Sánchez Alvarado, 2000). Because planarians belong to the Lophotrochozoan/Spiralian superclade, an animal grouping distinct from the Ecdysozoa (*e.g.*, flies and nematodes), yet share a common ancestor with the Deuterostomes (*e.g.*, vertebrates), *S. mediterranea* provides new opportunities to explore and inform the molecular processes underpinning the evolution of extant functional attributes of the animal epidermis.

Unlike the epidermis of adult flies and nematodes, the adult planarian epidermis is made from post-mitotic stem cell progeny that migrate from the internal mesenchyme, intercalate and then differentiate into the many different cell types that compose this organ, an observation originally made by Hallez in 1887 (Hallez, 1887). These migratory cells form part of a cohort of precursor epidermal cells that undergo multiple transition states on their way to epithelial differentiation (Tu et al., 2015). Lineage experiments have uncovered two populations of epidermal precursors. The first, defined by *Smed-prog-1*^*+*^ expression in early differentiating progeny; and the second characterized by the expression of *Smed-agat-1*^*+*^ in later, differentiating progeny (Eisenhoffer et al., 2008; van Wolfswinkel et al., 2014; Tu et al., 2015). Intriguingly, *in situ* hybridization of early and late progeny markers show little co-localization, suggesting that their transcriptional activation is spatiotemporally regulated (Zhu et al., 2015). Moreover, single-cell transcriptional profiling partitions the early and late progeny cells into two distinct populations (Wurtzel et al., 2015) and indicates that epidermal progenitors respond to positional cues along the planarian dorsoventral axis (Wurtzel et al., 2017). Together, these observations support the hypothesis that discrete changes must be occurring in the transcriptional landscape supporting epidermal lineage progression in planarians.

The intricate spatiotemporal differentiation of precursors into epidermal cells is reflected by the transcriptional complexity recently uncovered in planarian neoblasts. These studies identified a sub-population of neoblasts expressing *Smed-zfp-1* and *Smed-p53 (i.e.,* the zeta-class) to be the origin of epidermal progenitors (van Wolfswinkel et al., 2014). RNAi knockdown of *zfp-1* or *p53* reduces the number of intermediate precursors for the epidermal lineage. Such precursor depletion ultimately leads to ventral curling of the animals, a phenotype attributed to loss of epidermal integrity preferentially in the ventral epidermis (Pearson and Sánchez Alvarado, 2010; Wagner et al., 2012). Even more strikingly, *zfp-1* RNAi depletes the entire zeta-class population, yet leaves other classes of neoblasts *(i.e.,* the sigma-and gamma-classes) unaffected (van Wolfswinkel et al., 2014), suggesting that *zfp-1* functions specifically in the epidermal lineage. *Smed-zfp-1* encodes a zinc-finger protein with a highly conserved C2H2 DNA binding domains (Wagner et al., 2012), while *Smed-p53* shares a highly conserved DNA-binding domain with that of vertebrate *TP63* (Pearson and Sánchez Alvarado, 2010).

In this study, we dissect the transcriptional regulatory network underlying the planarian epidermal lineage during homeostasis. We describe several conserved transcription regulators and demonstrate a fundamental, upstream role for *p53* in planarians in the transcriptional cascade leading to epidermal lineage progression. Furthermore, we identify two transcription factors that together are not generally associated with the epidermal lineages of other animals: a sry-box homolog, *Smed-soxP-3,* and a pair-box homolog, *Smed-pax-2/5/8* acting downstream of *p53* and *zfp-1* and functioning cooperatively to specify the identity of early progeny cells. We show that *SoxP-3* and *pax-2/5/8* are required for the expression of a novel family of secreted proteins, PROG, which are specific to the epidermal lineage and likely reflect a planarian-specific adaptation. Moreover, we describe for the first time novel PROG+ secretory vesicles whose production depends on *SoxP-3* activity. Altogether, our work has uncovered a defined set of transcriptional modules integrated into a hierarchical regulatory program that likely drive the specification and homeostasis of the planarian epidermis, adding to our understanding of mechanisms underpinning epidermal cell diversity in animals.

## Results

### *p53-* and *zfp-l(RNAi)* yield similar transcriptional changes in planarian stem cells likely associated with epidermal differentiation

The zeta-class neoblasts associated with the epidermal lineage are characterized by their higher expression of *zfp-1, fgfr-1, p53, soxP-3,* and *egr-1* (van Wolfswinkel et al., 2014). However, only *zfp-1* and *p53* have been reported to induce a ventral curling phenotype after RNAi (Pearson and Sánchez Alvarado, 2010; Wagner et al., 2012). We sought to take advantage of these data to identify regulators of epidermal differentiation. We first confirmed that *zfp-1* and *p53* RNAi each depleted early and late progeny cells with similar kinetics (Figure 1 Supplement 1A, B). Second, we determined the extent of co-expression of these transcription factors *in vivo* in neoblasts (*piwi-l*^*+*^ cells) via fluorescent *in situ* hybridization (FISH) and showed that *zfp-1* and *p53* co-localized extensively to the *piwi-l*^*+*^ compartment (Figure 1 Supplement 1C), suggesting that they function in the same stem cell sub-population. *p53,* however, was detected in a broader domain spanning both neoblasts and post-mitotic progeny cells (*piwi-l*^*neg*^ prog-1^+^), indicating that *p53* may have multiple, cell lineage-specific roles (Figure Supplement 1C).

**Figure 1.**
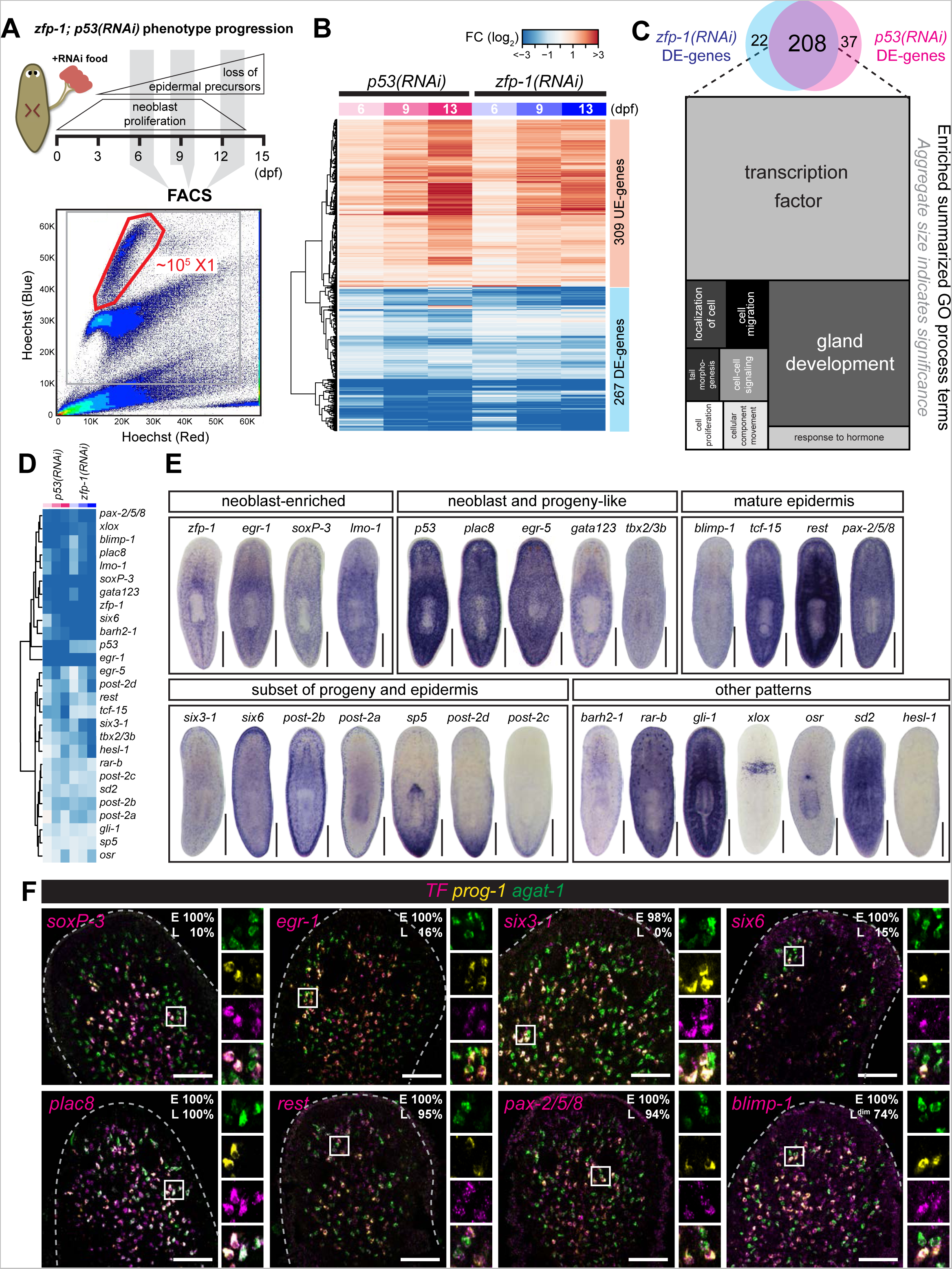
*p53-* and *zfp-l(RNAi)* analyses define shared transcriptional changes in neoblasts and their differentiation progeny. (A) Schematic of experimental strategy. Planarians fed with dsRNA food were dissociated into cell suspensions on 6, 9, and 13 dpf, and the neoblasts were FACS-sorted for RNA-Seq. A representative FACS plot (control 9 dpf) shows the gating of dividing neoblasts (XI). (B) RNA-Seq analysis of 576 genes shows highly correlated expression profiles in *p53-* and *zfp-l(RNAi)* X1 cells (selection criteria: FC > 1.5, p-adj < 0.01, ≥ 2 time points). Each column represents data of one time point. DE, down-regulated in expression. UE, up-regulated in expression. Colors represent log_2_ fold change (FC) to control. Color key is the same in Figure 1B and 1D. (C) Venn diagram of *p53-* and *zfp-l(RNAi)* DE-genes and the major GO terms enriched in the 208 common DE-genes. Related GO terms are merged in the same shade and the size indicates significance of a group of GO terms (Supek et al., 2011). (D) Expression profiles of the transcription factors (TFs) identified in the common DE-genes of *p53-* and *zfp-1 (RNAi)* X1 cells. (E) WISH patterns of candidate TFs. Scale bar, 500μm. (F) Co-localization of TF, *prog-1* and *agat-1* analyzed by FISH. Box region is split into individual channels to the right. Percentage of early (E) or late (L) progeny cells detected positive for candidate TF are shown in the upper right corner. Scale bar, 100μm.

We then profiled by RNA-Seq genome-wide transcriptional changes of neoblasts (“X1” population) isolated from *p53-* and *zfp-1 (RNAi)* animals to systematically identify potential regulators involved in epidermal differentiation at the neoblast stage (Figure 1A). It was reported that RNAi of *zfp-1* specifically eliminates the zeta-class neoblasts without affecting other progenitors that regenerate brain, muscle, gut and protonephridia (van Wolfswinkel et al., 2014). In contrast, RNAi of *p53* induces hyper-proliferation of neoblasts followed by rapid decline of these cells (Figure 1 Supplement 1D, F) (Pearson and Sánchez Alvarado, 2010). Based on the onset of the *zfp-1* and *p53* phenotypes, we chose to isolate X1 cells at 6, 9 and 13 dpf to include the earliest observed absence of early progeny cells, prior to the depletion of neoblasts in *p53(RN*Ai) animals (Figure 1A). We identified a total of 576 genes that showed changes in expression of at least 1.5 fold and an adjusted p-value < 0.01 for at least two time points in either *p53-* or *zfp-l(RNAi)* X1 populations (Figure 1B; Supplementary Table 1). The expression levels of these genes showed similar kinetics in *p53-* and *zfp-1 (RNAi)* X1 cells by RNA-Seq (correlation coefficient *r =* 0.9, Figure 1B) indicating that *p53* and *zfp-1* regulate similar biological processes in neoblasts. We surveyed the expression levels of previously reported genes and confirmed that all known zeta-class markers are down-regulated in *p53-* and *zfp-1 (RNAi)* (Figure 1 Supplement 1E). Markers of other lineages are either unchanged or are up-regulated. In particular, neuronal and protonephridial lineage markers showed increased expression along the time course suggesting an increase of other tissue progenitor cells at the expense of the epidermal lineage (Figure 1 Supplement 1E).

We identified 208 genes with significantly decreased expression levels (DE-genes) in both *p53-* and *zfp-1(RNAi)* X1 cells (FC > 1.5, p-adj < 0.01, ≥ 2 time points) and analyzed the resulting gene set by assigning gene ontology (GO) terms (Figure 1C, Supplementary Table 2). We found that transcription factor activity (G0:0003700) and gland development (G0:0048732) are the most enriched terms. Among the identified 208 DE-genes, 105 genes were previously classified by single-cell sequencing (SCS) technology (Wurtzel et al., 2015), and 83/105 (79%) genes were assigned to the epidermal lineages (Supplementary Table 1). We hypothesized from these data that the transcriptionally identified gene set are likely enriched in regulators of epidermal differentiation.

We tested this hypothesis by selecting 27 transcription factors (TFs) from the DE-gene list, (Figure 1D), and performed whole-mount *in situ* hybridization (WISH) (Figure 1E). The majority of genes tested showed expression patterns characteristic of epidermal lineage (Figure 1E). For instance, like *zfp-1,* the genes *egr-1, soxP-3,* and *lmo-1* appear to be highly expressed in neoblasts. Other genes like *plac8, egr-5, gata123* and *tbx2/3b* are expressed in both neoblasts and discrete sub-epidermal cells in a manner that is not dissimilar to *p53* expression. Additionally, *blimp-1, tcf-15, rest,* and *pax-2/5/8* are expressed in both discrete sub-epidermal cells and in mature epidermis. We also found several TFs expressed in a subset of mature epidermal cells. For example, *six3-1, six6, post-2b,* and *post-2a* are all enriched in cells located at the dorsoventral boundary, possibly marking adhesion gland cells. Another subset of genes represented by *sp5, post-2d* and *abdBa-2* exhibited graded expression along the anteroposterior axis. The rest of the genes examined showed strong expression in non-epidermal tissues.

We then used irradiation to rapidly and specifically eliminate neoblasts and, subsequently, epidermal progeny cells (Eisenhoffer et al., 2008), and checked the expression levels of the above TFs in an irradiation time-course by RNA-Seq (Figure 1 Supplement 2; Supplementary Table 3). We found that only those genes with expression enriched in neoblasts and progeny cells *(zfp-1, egr-1, soxP-3, lmo-1, p53, egr-5, gata123, tcf-15* and *six6*) were significantly down-regulated upon irradiation (Figure 1 Supplement 2). We then performed double FISH of TFs with early (*prog-1*) and late (*agat-1*) progeny markers (summary in Supplementary Table 4). Among all TFs examined, expression of *soxP-3, egr-1, six3-1,* and *six6* show substantial co-localization with the early progeny marker *prog-1,* but not late progeny marker *agat-1.* In contrast, *plac8, rest, pax-2/5/8,* and *blimp-1* extend their expression to late progeny and mature epidermis (Figure 1F). These expression patterns reflect on a complex temporal and spatial regulation of transcription as cells transition from early to late progeny cell stages.

### RNAi screen of *p53-* and *zfp-1* -modulated genes identifies *soxP-3* and *pax-2/5/8* as early transcriptional regulators in epidermal precursors

Based on their spatial distribution and association with progenitor cell markers (Figure 1F, Supplementary Table 4), we investigated 17 TFs for their possible roles in the differentiation of progeny cells. RNAi animals for each of the tested TFs were fixed and examined by WISH for *prog-1, prog-2* and *agat-1* expression (Figure 2A). Notably, of all tested TFs, only *soxP-3-* and *pax-2/5/8(RNAi)* animals showed significant reduction in expression of the early progeny markers *prog-1* and *prog-2,* but not the late progeny marker *agat-1.* This result was unexpected, as we and others have hypothesized that the late progeny arise from the differentiation of early progeny. Two possibilities may explain this result. First, that the early progeny cells are being eliminated in *soxP-3-* and *pax-2/5/8 (RNAi)* animals with the late progeny cells originating from another lineage that is independent from the early progeny cells. Second, the early progeny cells in *soxP-3-* and *pax-2/5/8 (RNAi)* animals are not expressing *prog-1 and prog-2,* markers but are still capable of differentiating into late progeny cells. To better understand the effects of *soxP-3-* and *pax-2/5/8(RNAi*) on the early stage of epidermal differentiation, we examined the expression of additional markers specific to the early progeny cells.

**Figure 2.**
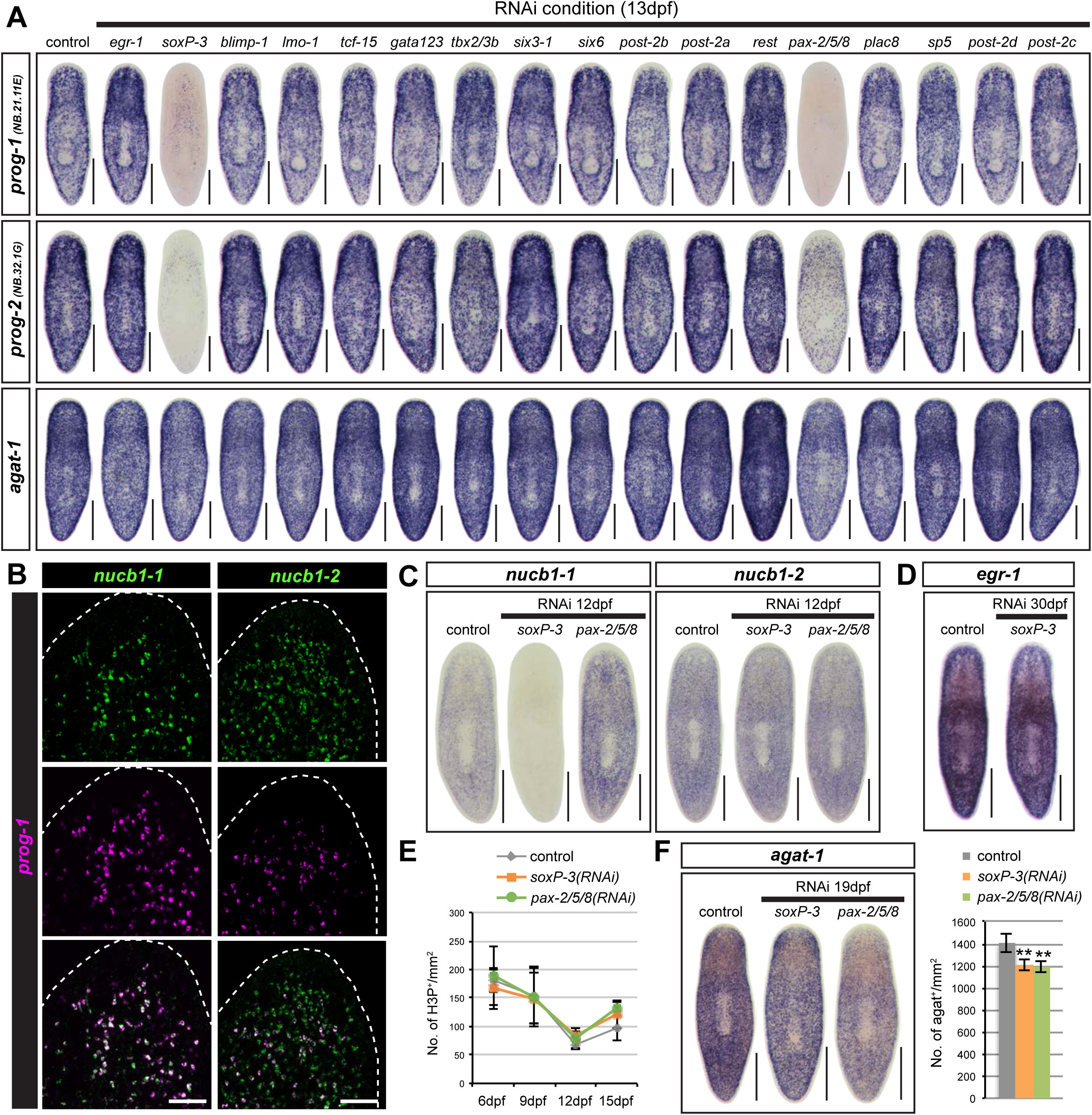
*soxP-3* and *pax-2/5/8* regulate early epidermal precursor trasncription. (A) *soxP-3-* and *pax-2/5/8(RNAi)* animals show reduced WISH signals of early progeny markers (*prog-1* and *prog-2,* 100% penetrance) but not late progeny marker *agat-1* in the TF RNAi screen (n > 8 per RNAi, 2 independent experiments). Scale bar, 500μm. (B) *nucbl-1* is 100% co-localized with *prog-1* analyzed by FISH, *nucbl-2* is partially overlapped with *prog-1* and may label late progeny cells. Scale bar, 100μm. (C) Expression of *nucbl-1* and *nucbl-2* analyzed by WISH, *nucbl-1* expression is eliminated in *soxP-3(RNAi)* animals but is up-regulated in *pax-2/5/8(RNAi)* animals (n > 6 per RNAi, 100% penetrance), *nucbl-2* expression is unaffected in both *soxP-3-* and *pax-2/5/8(RNAi)* animals. Scale bar, 500 μm. (D) *egr-1* expression is unaffected by *soxP-3* RNAi. Scale bar, 500μm. (E) Neoblast proliferation is unaffected by *soxP-3-* and *pax-2/5/8* RNAi analyzed by H3P immunostaining (mean ± s.d., n > 8 per time point). (F) *agat-1*^*+*^ cells are slightly reduced in *soxP-3-* and *pax-2/5/8(RNAi)* animals. Scale bar, 500μm. Quantification of *agat-1*^+^ cells is shown in the bar graph (mean ± s.d., n > 8). Double asterisks, p<0.001. Unpaired Student’s t-test.

Two *nucleobindin-1 (nucbl*) homologs, *nucbl-1* and *nucbl-2* were previously shown to be expressed in the epidermal lineage (Tu et al., 2015; Zhu et al., 2015). *nucb1-1* is expressed only in the early progeny cells whereas *nucb1-2* is expressed in both early and likely late progeny cells (Figure 2B). Like other early progeny markers in the *soxP-3 (RNAi), nucb1-1* expression is also undetectable. Surprisingly, *nucb1-1* expression level increases upon *pax-2/5/8* RNAi knockdown (Figure 3C) suggesting that early progeny cells are indeed still present in these animals and capable of expressing other early markers. Because *egr-1* expression is co-localized 100% with early progeny cells (Figure 2C), we looked at its expression in *soxP-3* RNAi animals and found no difference from control animals (Figure 3D). These results indicate that abrogation of both *soxP-3* and *pax-2/5/8* by RNAi affects the transcriptional output of the early progeny genes rather than the viability of the cells.

**Figure 3.**
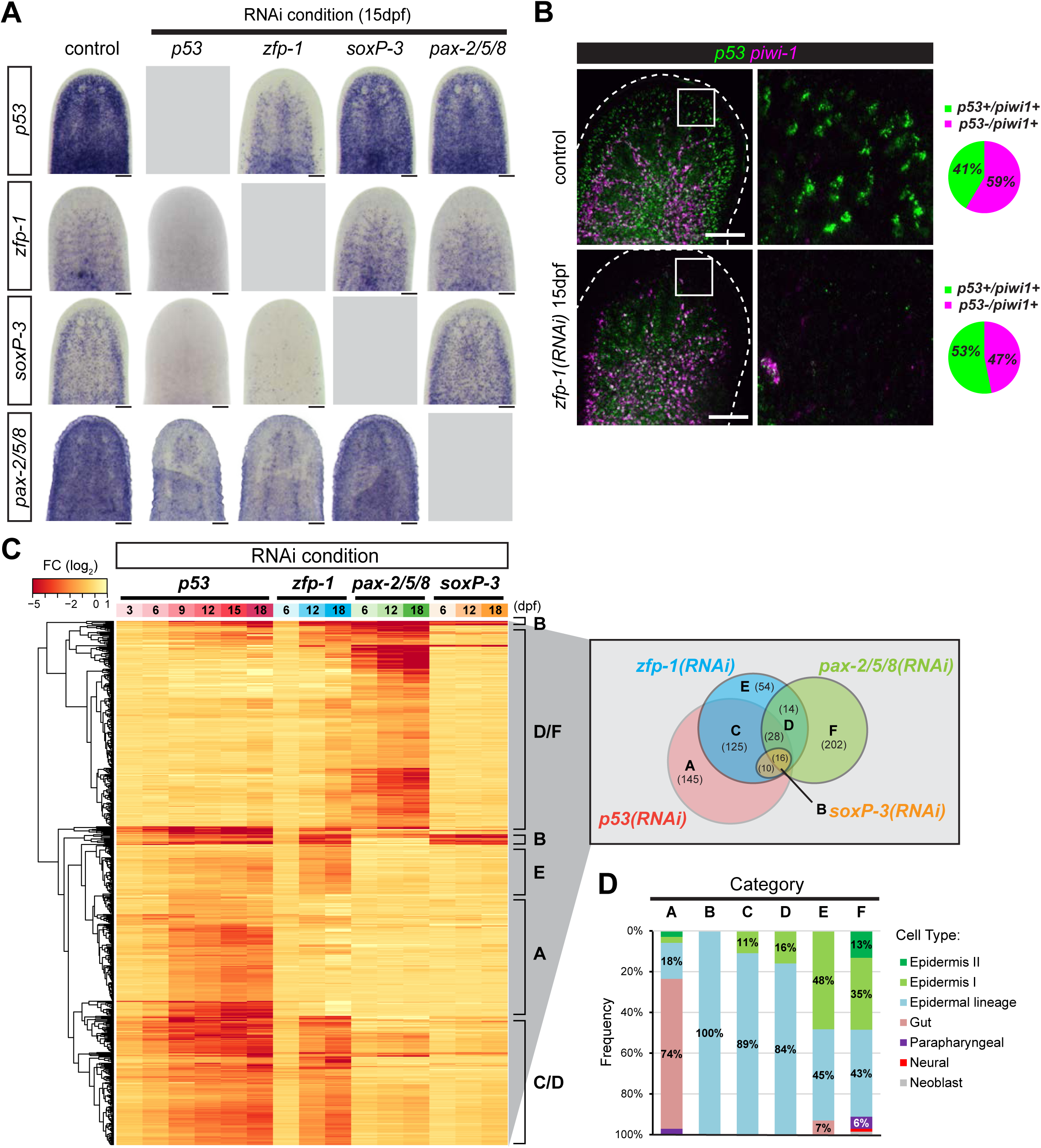
*p53, zfp-1, soxP-3* and *pax-2/5/8* regulate shared and distinct gene sets in a hierarchical transcriptional network. (A) WISH of *zfp-1, p53, soxP-3* and *pax-2/5/8* in the corresponding RNAi. *p53* RNAi eliminates the expression of all three TFs (n > 8, 100% penetrance), *zfp-1* RNAi eliminates *soxP-3* and *pax-2/5/8* expression, and reduce *p53* expression in subset of mesenchymal cells, likely due to the loss of early and late progeny cells. *soxP-3* and *pax-2/5/8* RNAi do not affect the expression of any of the other three TFs. Scale bar, 100μm. (B) *p53* and *piwi-1* double FISH shows that *p53* expression in the neoblast is not reduced by *zfp-1* RNAi. Scale bar, 100μm. Quantification of *p53*^*+*^ neoblasts of the representative images is shown in the pie charts (total cells counted: control = 1028; *zfp-1 (RNAi) =* 648). (C) FISH of pooled riboprobes of sigma-class (*soxP-1* and *soxP-2*), gamma-class (*hnf −4* and *gata456),* and zeta-class *(zfp-1, egr-1, soxP-3* and *fgfr-1*) is performed to identify neoblast sub-populations and their co-localization with *p53.* Boxed region is split into individual channels to the right. Arrowheads point to the triple positive cells. Scale bar, 100μm. (D) Expression profiles of 594 DE-genes identified in the RNA-Seq analysis of *p53-, zfp-1-, pax-2/5/8-* and *soxP-3(RNAi)* animals. Selection criteria of the gene list: FC > 2, p-adj < 0.001, ≥ 2 time points, except for *p53(RNAi)* in which FC ≥ 2, p-adj < 0.001, > 4 time points were used. DE-genes are classified into 7 major categories (A-F) and the numbers of each category are shown in the Venn diagram. (E) Cell type specific genes (Wurtzel et al., 2015) represented in the DE-gene Categories. Category A contains large numbers of gut markers. Categories B/C/D/E/F contain genes expressed along the epidermal lineage (including early progeny, late progeny and mature epidermis). More genes expressed in the mature epidermis are present in Category E and F.

To test whether the epidermal lineage of *soxP-3-* and *pax-2/5/8(RNAi)* animals can still differentiate properly, we examined the expression of the late progeny markers *nucb1-2* and found no difference as compared to control (Figure 3C). Similarly, other previously identified late differentiation markers (Tu et al., 2015; Zhu et al., 2015) are expressed normally in *soxP-3(RNAi)* animals (Figure 2 Supplement 1) confirming that these early precursor cells are still capable of differentiating into later stages despite of their failure to express some of the early progeny markers. Additionally, neither *soxP-3-* nor *pax-2/5/8(RNAi)* induce neoblast proliferation over the time course in treated animals (Figure 2E), suggesting that the integrity of the epidermis is not severely compromised and that a wound response does not appear to be triggered as in the case of other downstream lineage effectors such as *egr-5(RNAi)* (Tu et al., 2015). Nevertheless, we did observe a small but significant difference in the density of *agat-1*^*+*^ cells in *soxP-3-* and *pax-2/5/8(RNAi)* animals (Figure 2F) suggesting that subtle phenotypes associated with cell fate allocation may exist under these conditions.

### Hierarchically deployed transcriptional events drive epidermal fate decisions

We sought to define the most likely role *soxP-3* and *pax-2/5/8* may play in a transcriptional network controlling planarian epidermal differentiation. Therefore, we performed reciprocal RNAi followed by WISH of each TF to determine their most likely epistatic relationships to *p53* and *zfp-1* in the epidermal lineage (Figure 3A). As expected, *soxP-3* and *pax-2/5/8* expression in the mesenchyme were undetectable in the *p53-* and *zfp-1 (RNAi)* animals. Conversely, *p53* and *zfp-1* expression were not affected by RNAi of *soxP-3* and *pax-2/5/8.* Furthermore, the expression of *soxP-3* was not dependent on *pax-2/5/8* and vice versa. These observations strongly indicated that *soxP-3* and *pax-2/5/8* act in parallel to each other and that both are downstream of *p53* and *zfp-1,* likely initiating the expression of early progeny genes cooperatively in the epidermal lineage. Moreover, the epistatic interactions uncovered by our work appear to be epidermal-lineage specific. We noted that *pax-2/5/8* expression is also detected in a subset of neuronal cells (Figure 3 Supplement 1), and that such expression is not immediately affected by *p53-* and *zfp-1 (RNAi),* suggesting that *pax-2/5/8* expression in those tissues may be independent of *p53* and *zfp-1* functions.

Interestingly, while *zfp-1* expression is undetectable in the *p53(RNAi)* animals, *p53* expression is only partially eliminated by *zfp-1* RNAi (Figure 4A, B). For instance, *p53*^*+*^*piwi-l*^*neg*^ cells in front of the photoreceptors *(e.g.,* differentiating early and late progeny cells) completely disappear in the *zfp-1 (RNAi)* animals. However, *p53*^*+*^*piwi-l*^*+*^ cells are still present in the mesenchyme in the *zfp-1 (RNAi)* animals although with a slight increase in their proportion (53% versus 41% in control, Figure 3B). These data suggest that *p53* expression in the neoblast compartment may be independent of *zfp-1* function, and that there may be increased proliferation in the *p53*^*+*^ neoblast pool in response to the loss of the epidermal lineage.

**Figure 4.**
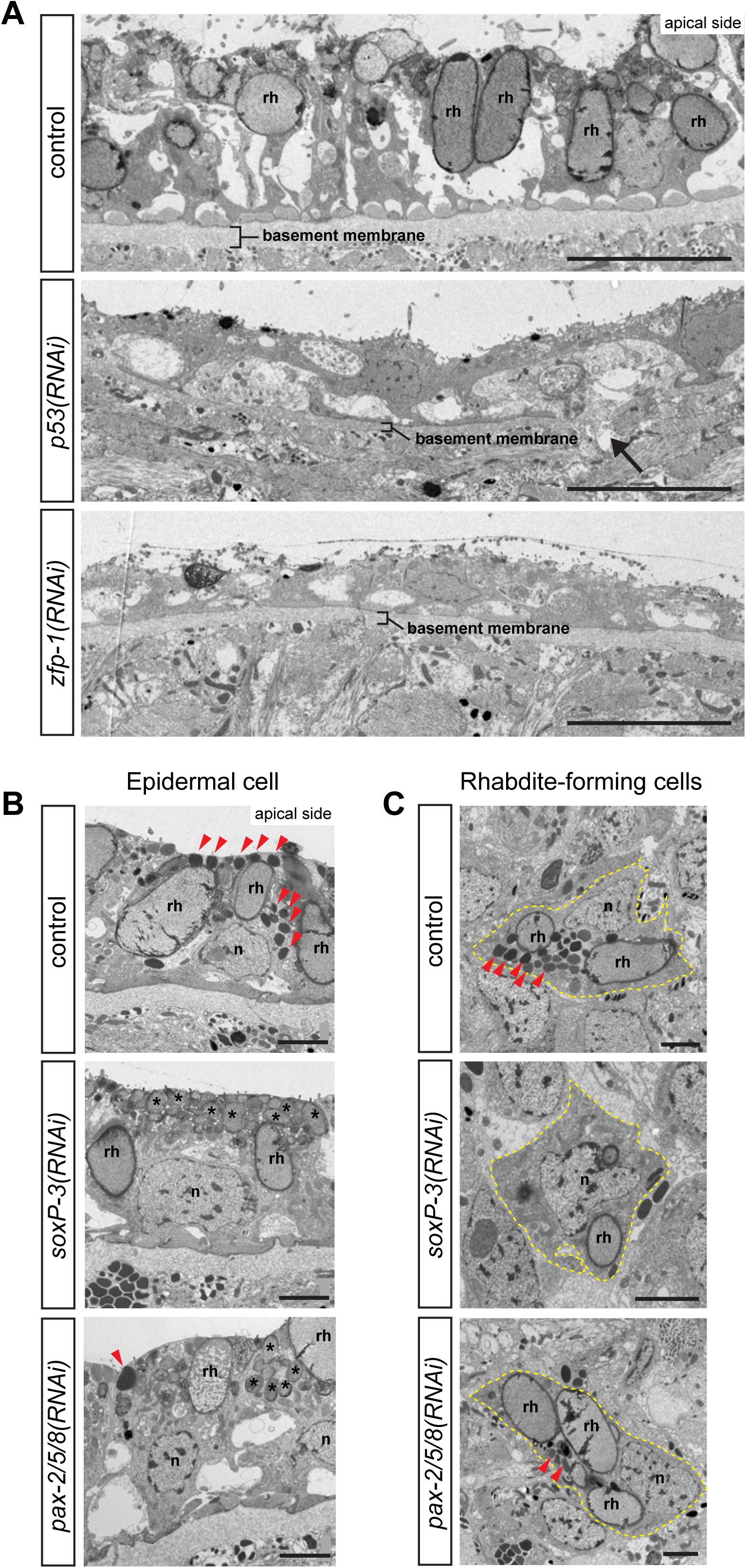
Ultrastructural analysis show distinct epidermal defects in *p53-, zfp-* 1-*, soxP-3-* and *pax-2/5/8 (RNAi)* animals. (A) TEM micrographs of ventral epidermis. *p53-* and *zfp-1 (RNAi)* epidermis appear to be stretched, contain reduced number of rhabdites (rh) and sit on thinner basement membrane (arrow points to a rupture), n = 3,100% penetrance. Scale bar, 10μm. (B) TEM micrographs of dorsal epidermis. The numbers of electron-dense granules (arrowheads) in the epidermal cells are greatly reduced by *soxP-3-* and *pax-2/5/8* RNAi. Instead, the apical side of *soxP-3* and *pax-2/5/8(RNAi)* epidermis are filled with rhabdite-like vesicles (asterisks), n, nucleus; rh, rhabdite. Scale bar, 2μm. (C) TEM micrographs of rhabdite-forming cells. Electron-dense granules (arrowheads) observed in the control cell are greatly reduced in *soxP-3-* and *pax-2/5/8(RNAi)* cells. Scale bar, 2μm.

### *p53, zfp-1, soxP-3* and *pax-2/5/8* regulate shared and distinct gene sets in the epidermal lineage

To investigate the functions that each of the identified TFs may play in the postmitotic differentiation of the epidermal lineage cells, we generated RNA-Seq datasets of *zfp-1-, soxP-3-* and *pax-2/5/8(RNAi)* animals on 6, 12, 18 dpf. Together with a previous *p53(RNAi)* dataset (Tu et al., 2015), we selected for further analysis 594 DE-genes significantly down-regulated by 2-fold with an adjusted p-value < 0.001 in at least 2 time points in the *zfp-1-, soxP-3-* or *pax-2/5/8(RNAi)* time courses, and at least 4 time points of the *p53(RNAi)* time course (Venn diagram in Figure 3C; Supplementary Table 5). Based on the expression profiles across different RNAi conditions, we catalogued these DE-genes into 6 major categories (A-F) (Figure 3C). 145 genes in Category A are specifically down-regulated in *p53(RNAi)* but not in other TF RNAi conditions. Category B (the smallest gene set) is composed of genes that are down-regulated in *soxP-3 (RNAi).* Categories C/D/E contain *zfp-1(RNAi)* DE-genes but may or may not be down-regulated in other RNAi conditions. And Category F encompasses genes specifically down-regulated in *pax-2/5/8(RNAi).*

We analyzed the data for the presence of cell types influenced by each TF by correlating the DE-genes of each category to a previously published list of cell-type specific markers (Wurtzel et al., 2015) (Figure 3D). We found that DE-genes in Categories B/C/D/E/F contain markers specifically expressed in the epidermal lineage, with more mature epidermal markers in Category E and F. Interestingly, Category A, the *p53(RNAi)* DE-genes, showed enrichment with gut markers suggesting a role for *p53* in the gut or gut progenitors. We tested this prediction by analyzing gut morphology via TEM and observed severe defects in this tissue in *p53(RNAi)* animals but not in *zfp-1 (RNAi)* specimens (arrowhead, Figure 3 Supplement 2).

Taken together, the data indicate that the RNAi-defined epistasis (Figure 3A) and the corresponding analyses of global transcriptomic changes (Figure 3C) define an epidermal lineage progression that includes a neoblast stage in which the most upstream regulator is *p53,* which besides targeting genes important for the gut lineage (Figure 3 Supplement 2), also robustly drives the expression of *zfp-1* (Figure 3A). In turn, *zfp-1* appears to act cooperatively with *p53* to induce the expression of multiple TFs, including *soxP-3* and *pax-2/5/8* (Figure 3A and C), likely responsible for the maturation and diversification of the epidermal lineage (Figure 3D).

### Ultrastructural analyses implicate Rhabdite-forming cells are migratory epidermal precursors

In contrast to the readily detectable consequences of abrogating *p53-* and *zfp-1, soxP-3-* and *pax-2/5/8(RNAi)* failed to produce overt epidermal deficiencies. We were struck by the range of phenotypic severity caused by the RNAi of these 4 transcription factors on the planarian epidermis. To better understand these observations, we sought to take advantage of the precise spatial and temporal transcriptional regulation of the epidermal lineage to define ultrastructural characteristics associated with the specification and differentiation of epidermal cells. To guide our analyses, we focused our efforts on rhabdite-forming cells. Rhabdites are specialized, rod-shaped structures found in the epidermis of all flatworms and some nemertene worms (Walker and Anderson, 2001). Rhabdites are discharged in mucous secretions and have been associated with both defensive mechanisms (Hyman, 1951) and adhesion (Martin, 1978).

Rhabdites have distinct morphology and are relatively easy to locate and identify by TEM. We examined tiled TEM images of animal cross sections for the presence of both epidermal rhabdites and rhabdite-forming cells residing in the internal mesenchyme. Rhabdites were readily identified in both epidermal and mesenchymal cells in control animals (rh in Figure 4A, B), but were undetectable in either of these cell types in *p53-* and *zfp-1 (RNAi)* (Figure 4A and Figure 4 Supplement 1). Moreover, the epidermis of *p53-* and *zfp-1 (RNAi)* animals appeared stretched and deformed, and the basement membranes noticeably thinner (Figure 4A). Interestingly, we also noticed that while *p53(RNAi)* affected the presence of rhabdites in both dorsal and ventral epithelia, *zfp-1 (RNAi)* appeared to primarily affect ventral rhabdites (Figure 4 Supplement 1) supporting recent observations that epidermal progenitors respond to positional cues along the planarian dorsoventral axis (Wurtzel et al., 2017). These observations suggest that rhabdite-forming cells may be epidermal precursors and that in their absence epidermis homeostasis is severely compromised. In contrast, ultrastructural analyses of *pax-2/5/8-* and *soxP-3(RNAi)* animals revealed a marked reduction or a complete absence, respectively, of a cohort of dark granules normally found in both epidermal and rhabdite forming cells without noticeably altering the rhabdites (Figure 4B and C). Additionally, we found that the space at the apical side of the epidermal cell is instead filled with rhabdite-looking granules in the *soxP-3-* and *pax-2/5/8*fianimals. Altogether, we conclude from these data that planarian epidermal differentiation is likely a multi-step process in which neoblasts produce post-mitotic intermediate cells that progressively acquire features of mature epidermal cells (*e.g.,* rhabdites and other secreted granules) as they integrate into the epidermis.

### *soxP-3* and *pax-2/5/8* function cooperatively to regulate the expression of novel planarian genes

We noted that *soxP-3(RNAi)* specifically down-regulated a small number of genes (Figure 3C). Except for the homologs of nucleobindin (*nucb1-1*) and synaptotagmins (*syt-2* and *syt-14L),* all *soxP-3-*specific transcriptional changes are associated with novel genes devoid of clear homologs in other animal species. Some of these genes were previously identified by irradiation or FACS isolation of X1, X2 and Xin cell populations and named after *prog* (progeny) or *pmp* (post-mitotic progeny) (Eisenhoffer et al., 2008; Zhou et al., 2009) We examined the expression profiles of 21 *soxP-3(RNAi)* DE-genes identified via RNA-Seq from an irradiation time-course and found that they exhibit similar expression kinetics to *prog-1* and *prog-2* (Figure 5A) suggesting that they are expressed in the irradiation-sensitive early progeny cells. Indeed, double FISH labeling showed 100% co-localization between *prog-1* and the tested novel *prog* genes (Figure 5 Supplement 1A; (Zhu et al., 2015)). Interestingly, only 13 out of 21 *soxP-3(RNAi)* DE-genes were also down-regulated by *pax-2/5/8* RNAi (Figure 5A). We confirmed this by WISH and observed that while *prog-1, prog-1-1, prog-2* and *prog-2-5* expression levels are markedly reduced in *pax-2/5/8(RNAi)* animals, *prog-2-4* is only reduced moderately and *prog-2-2* expression is not noticeably affected when compared to controls (Figure 5 Supplement 1B). Since *soxP-3* and *pax-2/5/8* are expressed in all early progeny cells, we suspected that the organization of the *soxP-3(RNAi)* DE-genes may share common genomic features such as nearby DNA sequences that may correlate with their responsiveness to *soxP-3* and *pax-2/5/8* activities.

**Figure 5.**
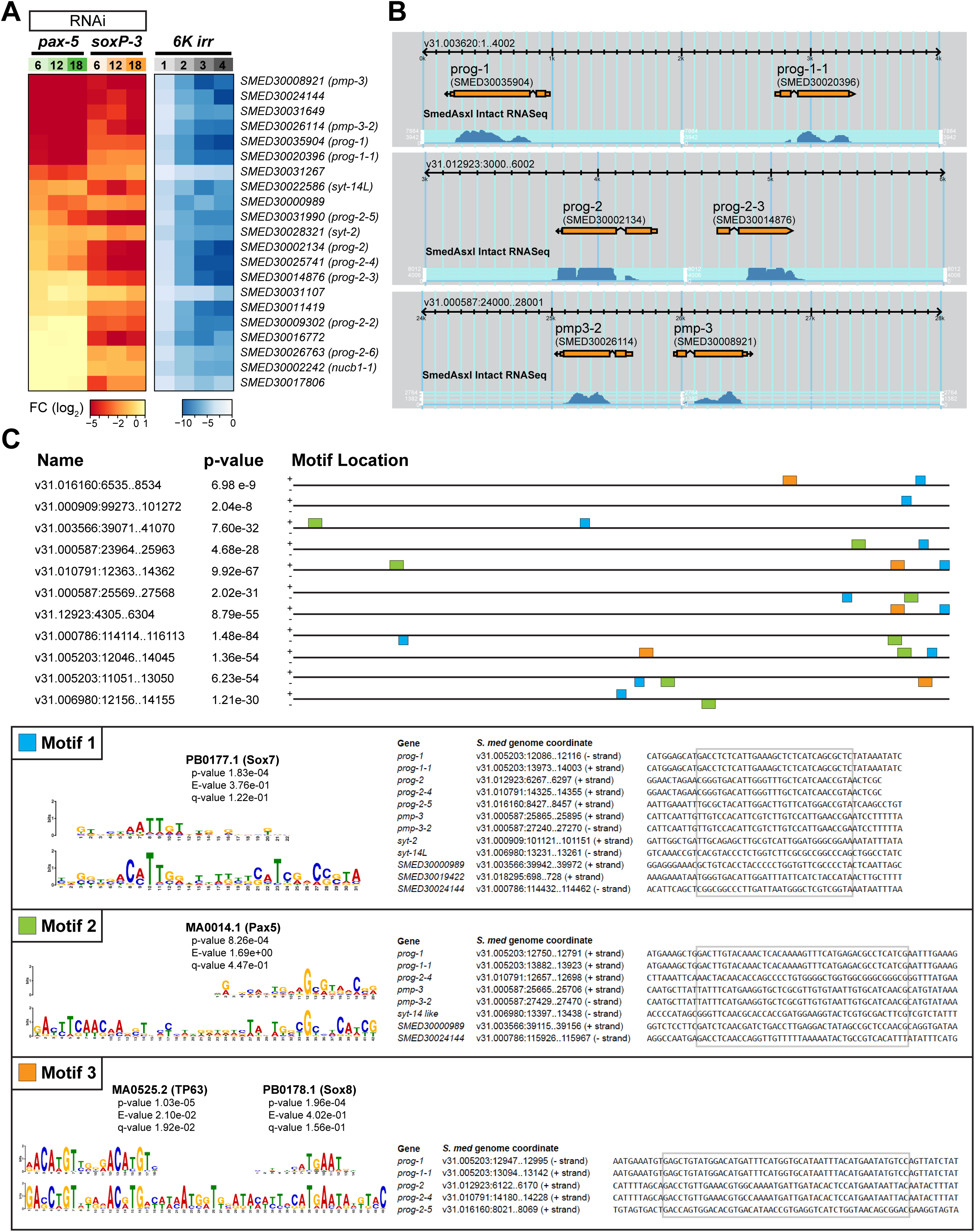
*soxP-3* and *pax-2/5/8* regulate the expression of a novel gene family uniquely arranged in the *S. mediterranea* genome. (A) Expression profiles of Category B DE-genes (from Figure 4D) in the *pax-2/5/8(RNAi), soxP-3(RNAi)* and irradiation time course RNA-Seq analysis. (B) Screenshot of SmedGD browser showing paired *prog* and *pmp* gene loci in the *S. mediterranea* genome V3.1 (Robb et al., 2015). (C) Top three motifs over-represented in the upstream 2kb sequences of DE-genes common in *soxP-3-* and *pax-2/5/8(RNAi).* The consensus matrix of vertebrate Sox7, Pax5, TP63 and Sox8 binding sites derived from TRANSFAC database and the genome coordinates of the motif sites are shown in the box.

We mapped the 13 DE-genes that are down-regulated in both *soxP-3-* and *pax-2/5/8(RNAi)* to the *S. mediterranea* genome V3.1 and obtained 2kb upstream of coding sequences for further analysis. Surprisingly, we found that some of these DE-genes are located in close proximity to each other and form opposing oriented “pairs” on the same genomic contig suggesting that they may share regulatory elements for transcription (Figure 5B Supplement 1). We performed *de novo* motif analysis using the MEME suite (Bailey et al., 2015) on 2kb upstream sequences from 64,028 mapable genes in the whole genome as background (Figure 5C). After we retrieved the top-enriched motifs, we compare the sequences to the JASPER database and identified at least three highly conserved TF binding sites (Gupta et al., 2007). Motif 1 is shared by 11 out of 13 DE-genes and contains a vertebrate Sox7 binding site. Motif 2 is identified in the eight DE-genes with higher fold-change in the *pax-2/5/8(RNAi)* animals including *prog-1, prog-1-1, pmp-3, pmp-3-2, SMED30000989* and *SMED30024144* and contains a vertebrate Pax-2/5/8 binding site. Motif 3 is found in the most highly expressed five DE-genes including *prog-1, prog-1-1, prog-2, prog-2-4* and *prog-2-5* (mean RPKM = 372 as compared to mean RPKM = 8.9 for the 8 remaining genes). Remarkably, Motif-3 contains not only a vertebrate Sox8 binding site, but also a highly conserved vertebrate TP63 binding site. We also find putative TP63 binding sites in the promoter region of *zfp-1* (Figure 5 Supplemental 2) Based on the highly homologous DNA-binding domains of planarian p53 and TP63, their binding sites may also be conserved. We conclude from these data that planarian p53 may bind to the identified putative regulatory elements and potentially modulate the subsequent binding of SoxP-3 and the resulting expression of *prog* genes.

### A novel gene family likely encodes epidermal secreted molecules in planarians

To further investigate the properties and functions of the novel *prog* genes, we performed SMART domain analysis (Schultz et al., 1998). We identified signal peptides at the N-terminus suggesting that the gene products may be secreted (Figure 6 Supplement 1A). In addition, the predicted proteins from these novel genes share similar architecture (coiled-coil and internal repeats) and nucleotide sequences. For example, *prog-2-2, prog-2-3, prog-2-4* and *prog-2-5* share higher than 50% identity at the nucleotide level (Zhu et al., 2015) suggesting that gene duplication events may have been involved in the evolution of this planarian-specific gene family.

**Figure 6.**
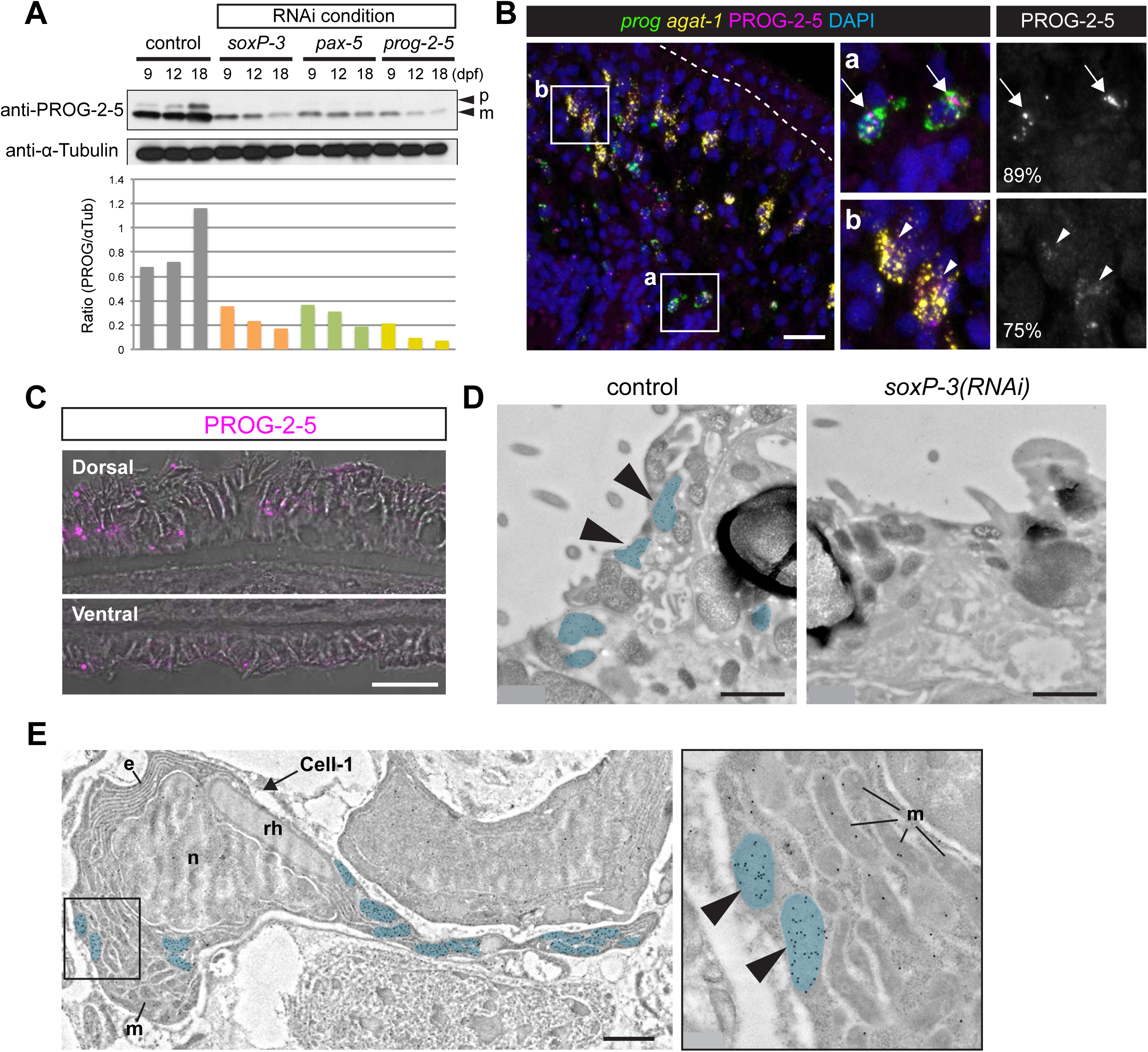
PROG-2-5 proteins are found associated with granules in the planarian epidermis. (A) Western blot analysis of PROG-2-5 in *soxP-3-, pax-2/5/8-* and *prog-2-5(RNAi)* whole worm lysates. Two bands corresponding to the precursor form (p) and mature form (m) of PROG-2-5 proteins are both significantly down-regulated in the RNAi lysates. Relative densities of PROG-2-5 bands to α-Tubulin are shown at the bottom. (B) Immunostaining of anti-PROG-2-5 combined with double FISH of *prog* (*prog-1 + prog-2* mix) and *agat-1.* Higher magnification of the boxed regions (a and b) is shown in the right panel. Large PROG-2-5+ puncta are observed in 89% of early progeny (*prog\**) cells (a, arrows). Smaller PROG-2-5^+^ puncta are also detected in 75% of late progeny (*agat-1*^*+*^) cells (b, arrowheads). Total 341 cells sampled from 5 animals are analyzed. Scale bar, 20μm. (C) Confocal/DIC images of PROG-2-5 immunostaining in the epidermis. Anti-PROG-2-5 detects puncta on both dorsal and ventral epidermis. Scale bar, 20μm. (D) IEM images showing the apical side of epidermal cells. In the control epidermal cell, immuno-gold labeled anti-PROG-2-5 antibodies detect signals in the oval-shaped granules (pseudo-colored in blue). PROG-2-5+ granules are undetectable in *soxP-3(RNAi)* epidermal cells, n = 2. Scale bar, μ.m. (E) IEM image of a mesenchymal cell labeled with anti-PROG-2-5 immuno-gold. Large rhod-shaped granules resembling rhabdites are found in the same cell labeled with PROG-2-5^+^ oval-shaped granules (pseudo-colored in blue) (also see Figure 7 -figure supplement 3). Higher magnification of the boxed region is shown in the right panel, e, endoplasmic reticulum; m, mitochondria; n, nucleus; rh, rhabdite. Scale bar, lμm.

We sought to investigate the cellular distribution of the novel PROG proteins by raising polyclonal antibodies against the protein product of *prog-2-5.* Purified polyclonal antibodies from two rabbits were tested for specificity on Western blots (Figure 6 Supplement 1B). The polyclonal obtained from Rabbit-2 not only displayed robust and specific reactivity, but also reproducibly detected two protein bands near 15 kDa standard. The two sizes likely correspond to the predicted full-length peptide (16.8 kDa) and the mature form of secreted products lacking the signal peptide (13.6 kDa). To verify specificity, we tested the antibodies on proteins obtained from *soxP-3-, pax-2/5/8-,* and *prog-2-5(RNAi)* animals and observed that the two different sizes of PROG-2-5 were both decreased in relation to control (Figure 6A). We conclude from these data that the obtained polyclonal antibodies are likely specific to *prog-2-5* protein products.

The newly developed polyclonal antibodies revealed that the mature form of PROG-2-5 (13.6 kDa) was detectable in *p53(RNAi)* samples even after 20 dpf when all *prog-2-5* transcript expressing early progeny cells are gone (Figure 6 Supplement 1C). We interpreted this result by hypothesizing that the protein may have a longer half-life than the mRNA that codifies it. As such, we would expect mature PROG-2-5 proteins to be distributed in places other than the early progeny cells. We tested this hypothesis by developing immunofluorescence (IF) assays for anti-PROG-2-5 in combination with FISH of epidermal lineage markers. We observed a 89% of co-localization of PROG-2-5 puncta with early progeny cells confirming the specificity of the IF staining (Figure 6B, Figure 6 Supplement 2). In addition to the bright puncta observed in the early progeny cells, we also detected less well defined PROG-2-5 IF signal in the majority of the late progeny cells (Figure 6B), and in the mature epidermis on both the dorsal and ventral surface of the animals (Figure 6C, Figure 6 Supplement 3). We conclude from these data that PROG-2-5 proteins are first produced in the epidermal progenitor cells as they differentiate and mature.

### PROG-2-5 localizes to discrete vesicles in both precursor and mature epidermal cells

The punctate staining pattern of PROG-2-5 suggested that this protein may be associated with specific organelles such as the rhabdites or the many vesicles found in the cells of the epidermal lineage (Figure 4B and C). In order to examine the subcellular localization of PROG-2-5, we developed reagents and protocols for immune-electron-microscopy (IEM) detection of proteins on fixed planarian sections. The immuno-gold-labeled anti-PROG-2-5 antibodies detected 0.5-μ.m-size granules not only on the epidermis (Figure 6D), but also in a cohort of mesenchymal cells with quite distinctive ultrastructural morphology, namely, elongated cell processes, an abundance of mitochondria and secretory granules, densely packed granular ER around the nucleus, and large rod-shape granules (rhabdite precursors) (Figure 6E, Figure 6 Supplement 3). These data clearly indicated that the vesicles containing PROG-2-5 immunogold label are distinct from rhabdites in both precursor and mature cells. Importantly, we find that *soxP-3(RNAi)* abolishes the production of the PROG-2-5 containing vesicles (Figure 6D, Figure 4B and C). Therefore, we conclude from these data that PROG-2-5 localizes to a distinct and previously undescribed secreted organelle whose production is dependent on the transcriptional activity of *soxP-3.* We refer to this organelle as the Hyman vesicle (see discussion for details).

## Discussion

The homeostatic dynamics and cellular diversity of the planarian epidermis provides an experimentally accessible model to study fundamental aspects of stem cell fate regulation and developmental biology. Here, we uncovered striking molecular similarities underpinning the epidermal lineage progression programs of planarians and mammals. Further, we identified novel components unique to planarians both at the regulatory network and gene level. Together with the phylogenetic position of planarians in relation to both vertebrates and traditional invertebrate model systems *(e.g.,* flies and nematodes), the planarian epidermis provides unique opportunities to inform our understanding of how conserved and derived mechanisms may have evolved to produce, maintain and diversify the metazoan epidermis.

### Epidermal lineage progression is mediated by a hierarchical transcriptional program

The planarian epidermal lineage is thought to transition through several morphologically and molecularly distinct stages before reaching the body surface (van Wolfswinkel et al., 2014; Tu et al., 2015). Here, we demonstrate that P53 likely initiates a hierarchical transcriptional program mediated by a defined cohort of TFs that sequentially activate gene expression as neoblast progressively differentiate into epidermal cells (Figure 7C). Our RNAi data support a model in which *zfp-1* is a transcriptional target of P53. Indeed, we found putative, highly conserved, vertebrate-like TP63 binding sites within a 5kb DNA region upstream the likely transcription start site of the *zfp-1* locus (Figure 5 Supplement 2; Supplementary Table 7). ZFP-1, in turn, likely acts cooperatively with P53 to control the expression of other downstream TFs associated with epidermal functions (Figure 7C). This feed-forward, hierarchical arrangement of epidermal-associated TFs in planarians is characteristic of gene regulatory networks that mediate developmental processes, and often induce the sequential expression of gene repertoires that effect the functional and morphological changes of a cell (Erwin and Davidson, 2009).

**Figure 7.**
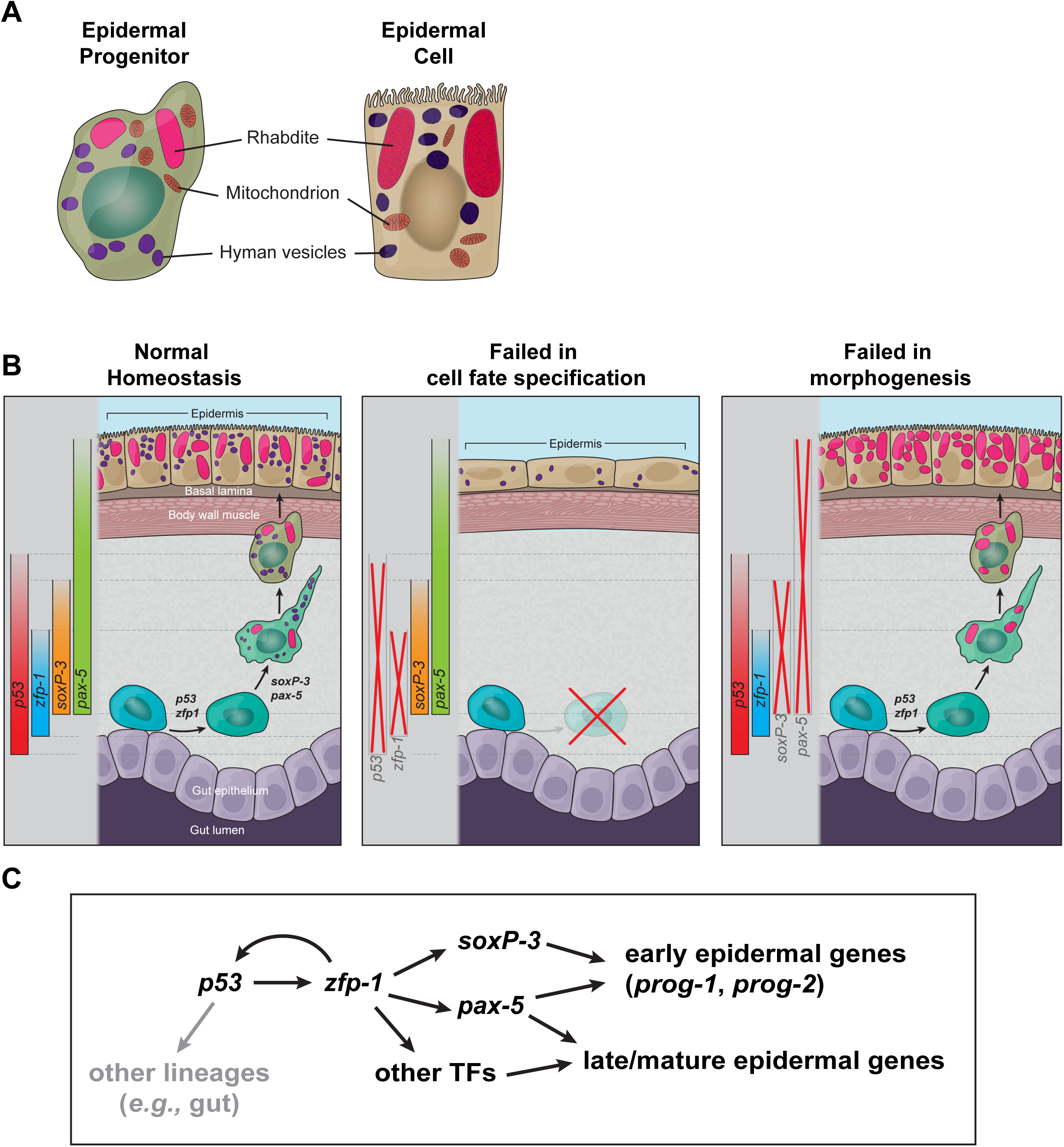
Model of epidermal lineage progression in planarian. (A) Schematic of ultrastructures of epidermal precursor and mature epidermal cell. Epidermal precursors contain features of the mature epidermal cells including large rod-shape rhabdites and small oval-shape granules. They are also abundant in mitochondria. (B) Schematic of the planarian epidermal lineage progression and the phenotypic differences in *p53-, zfp-1-, soxP-3* and *pax-2/5/8(RNAi)* animals. (C) Proposed transcription network in the epidermal lineage progression. Gray arrow represents aspects of the network not studied in depth.

The observed spatiotemporal transcriptional organization of the planarian epidermal lineage may be a consequence of transcriptional priming, an epigenetic mechanism thought to act at enhancers to bookmark loci for transcriptional activity at later stages of differentiation (Vernimmen and Bickmore, 2015). In fact, TP63 is thought to function as a pioneer factor, and binding of TP63 to chromatin is associated with the deposition of transcriptional activation marks that provide instructive cues that facilitate the binding of other TFs (Kouwenhoven et al., 2015). Given the recent demonstration that *p63* depletion in mice results in significant changes in the expression of genes involved in chromatin organization, remodeling and epigenetic modification (Boxer et al., 2014; Sun et al., 2015), it should be fruitful to determine whether SMEDP53 functions as a pioneer factor or not. In addition to *zfp-1,* the identification of predicted and highly conserved TP63 binding sites in some of the early progeny genes suggests that planarian P53 may facilitate SOXP-3 and PAX-2/5/8 binding. We have recently established an experimental approach for quantitatively assessing epigenetic landscapes in planarian neoblasts (Duncan et al., 2015) that could be extended to measuring transcription factor occupancy via chromatin immunoprecipitation (ChIP). Efforts to develop a ChIP-grade antibody against planarian P53 are currently underway as this will be an essential reagent in understanding not only how p53 influences the activity of downstream TFs, but also whether it too plays a role in regulating the global epigenetic landscape of the epidermal progenitor cells.

### The planarian epidermis as an invertebrate model to study dynamic epidermal organs

The regulatory association of *Smed-p53* and *zfp-1* is reminiscent of what is presently known about mammalian epidermis homeostasis, where one of the key targets of P63 is also a zinc finger protein, *i.e.,* ZNF750 (Sen et al., 2012; Zarnegar et al., 2012). Even though, *zfp-1* and *znf750* are both members of the large family of zinc-finger proteins, their functional homology remains to be established. However, we find it curious that a network architecture involving members of two distinct family of proteins (TP53/63 and ZNF) share similar gene regulatory architecture and hierarchy in both planarian and mammalian epidermal development. The P63/ZNF750 network in the epidermis has been a focus of intense research for its relevance to many human pathologies including atopic dermatitis, psoriasis, ichthyosis vulgaris, and skin cancer (Lopez-Pajares et al., 2013). However, because of the complexity of p63 isoforms and the crosstalk between p53/p73 and p63 due to their highly conserved DNA-binding domains, it has been a challenge to dissect p63 mediated downstream pathways responsible for its diverse functions (Vanbokhoven et al., 2011). Additionally, it is not known whether deletion of ZNF750 will cause developmental defects in epidermal tissue other than psoriasis, which is caused by point mutations that lead to hypomorph phenotype of this gene in human patients, as there are presently no published reports on the targeted elimination of this gene in mice. Nevertheless, based on the data presented here, the fact that the only member of the P53 family of proteins in *S. mediterranea* is P53, and that the *p53* locus only encodes a single mRNA species (Pearson and Sánchez Alvarado, 2010), we propose that planarians offer a unique opportunity to resolve the spatiotemporal transcriptional organization of dynamic epithelia. If in fact such network architecture shares a common evolutionary origin, a clear and testable prediction would be that abrogation of ZNF750 in the mouse epidermis should result in early lineage epidermal deficiencies similar to those observed in *zfp-1(RNAi)* treated planarians.

Altogether, our data establish an intriguing parallel between mammals and planarians of key molecular mechanisms driving the self-renewing capacity and commitment of progenitor cells in the respective epidermal tissues of these organisms.

### p53 family may play an ancient role in generating ectodermally derived tissues

In vertebrates, the *p53* gene family of transcription factors is composed of three members (*tp53*, *tp63* and *tp73*) and is known to play essential roles in tumor suppression, development and germ-line protection (Dötsch et al., 2010). Phylogenetic analyses of vertebrate *p53* family members suggested that they originated from a *tp63-like* common ancestor (Joerger et al., 2014). Our study supports this finding. We show that *Smedp53* is essential for maintaining the homeostasis of the planarian epidermis, a role played in the mammalian epidermis by the vertebrate *TP63.* This observation suggests that an ancient role of a P53 family member in epidermal development may have predated the split of deuterostomes and protostomes. How this genetic program may have been lost in flies and *C. elegans* is unknown. One explanation might be that the mature adult epidermis of both organisms does not turn over as rapidly as the vertebrate stratified epidermis or the planarian epidermis. Therefore, flies and nematodes likely evolved distinct developmental strategies by deploying alternative genetic programs. This is supported by recent studies of adult epidermal wound healing and repair in the fly *D. melanogaster.* In this organism, repair of damaged abdominal epidermis does not involve the recruitment and/or activation of epithelial progenitors, but rather the polyploidization of the surviving epidermal cells surrounding the wound via activation of Hippo and JNK signaling, resulting in efficient covering and healing of the damaged epidermis (Losick et al., 2013; Losick et al., 2016). Studies of more clades of animals that renew and regenerate their epidermis will be necessary to further investigate this hypothesis.

### A conserved Sox/Pax transcription factor pair is deployed to drive specific aspects of planarian epidermal morphogenesis

The Pax and Sox families of transcription factors are known for their diverse functions in development and disease. In the vertebrate eye development, Sox2 partners with Pax6 to activate the expression of δ-crystallin, a water-soluble structural protein found in the lens and the cornea (Kamachi et al., 2001). Interestingly, Pax-6 alone binds poorly to the lens-specific enhancer DC5 and requires the binding of Sox-2 to DC5 to facilitate the formation of a ternary complex of Pax6/Sox2/DC5 (Narasimhan et al., 2015). Remarkably, the reporter carrying vertebrate DC5 enhancer is activated by Drosophila Pax and Sox homologs, D-Pax2 and SoxN, in the Crystallin secreting cone cells of the compound eyes suggesting that Pax/Sox cooperative binding is evolutionary conserved.

In the epidermis, SOX and PAX proteins appear to act independently of each other. The SOX family of transcription factors, for example, has been extensively implicated in the modulation of fate lineages in the mammalian epidermis, particularly in driving the differentiation of epidermal fibroblasts into dermal papillae and keratinocytes (Driskell et al., 2009; Driskell et al., 2013; Lesko et al., 2013; Perdigoto et al., 2014). In addition, the highly specialized mechanoreceptor of the vertebrate skin develop thorough an entirely postmitotic and progressive maturation process that is also specified by SOX2 (Perdigoto et al., 2014). Interestingly, a role for PAX6 proteins in the maturation of Merkel cells has been also identified and appears to be the only reported function of this transcription factor in epidermal development and maintenance thus far (Parisi and Collinson, 2012).

Our study, however, found a similar role for a member of the PAX family of transcription factors in modulating the fate of epidermal progenitors (Figures 2A and 3A). To our knowledge, a partnership of Sox/Pax proteins in the epidermis has not yet been described. Our data indicate that planarian *soxP-3* and *pax-2/5/8* function cooperatively in the expression of a subset of *prog* genes during epidermal differentiation. Given their genomic architecture (Figure 5B), the newly identified *prog* genes are likely the result of gene duplication in the *S. mediterranea* genome, and may be a recently evolved gene family for an adaptation specific to planarians. Irrespective of the evolutionary origin of the PROG family of proteins, it remains intriguing that their regulation appears to be under the control PAX and SOX proteins, which themselves belong to gene families that can be dated back to the appearance of multicellularity (Degnan et al., 2009). Therefore, the SoxP-3/Pax-2/5/8 pair controlling the expression of *prog* genes may be a re-deployment of another more ancient network, which was integrated into the P53-mediated epidermal program. Given the roles that the TP53, SOX and PAX families of proteins play in the maintenance, diversification and specialization of the mammalian and planarian epidermis, *S. mediterranea* provides us with an opportunity to readily explore how conserved gene regulatory networks may be co-opted to drive the emergence of new specialized structures in evolutionarily conserved tissues.

### Rhabdite-forming cells are epidermal precursors in planarians

Rhabdites are large rod-shaped granules found in the epidermis of most Turbellarians (Hyman, 1951). They are embedded in the epidermis and are discharged when the animals are irritated. Planarian rhabdites are thought to contain mucopolysaccharides that form a mucous cuticle protecting the animal from environmental chemicals and pathogens. They are also deposited in the slimy tracks left behind by the animals, and therefore are also suggested to facilitate cilia-based gliding (Martin, 1978; Rompolas et al., 2010). Although the function of rhabdites have not been formally tested, this epidermal specialization is likely to play important roles in the interaction between planarians and their environment.

Previous studies have documented that rhabdite-forming cells in the mesenchyme may be epidermal precursors in some triclad species. Skaer observed that rhabdite-forming cells develop at the later stage of the embryogenesis of *Polycelis tenuis* embryos, and become the animals’ mature epidermis before they hatch (Skaer, 1965). Hori observed that rhabdite-forming cells participate in the regeneration of epidermis in the amputated *Dugesia japonica* (Hori, 1978). Here we made polyclonal antibodies that successfully detect the expression of an early progeny marker, PROG-2-5, and observed co-localization of PROG-2-5 with rhabdite-forming cells in the planarian *S. mediterranea.* Moreover, *p53* and *zfp-1* RNAi eliminate rhabdite-forming cells in the mesenchyme supporting that they are epidermal precursors (Figure 4 Supplement 1). Our molecular and genetic studies, therefore, unambiguously validate previous ultrastructural and morphological observations, and confirm that these otherwise mysterious mesenchymal cells with distinct organelles are indeed planarian epidermal precursors. We postulate that the combined activities of *p53* and *zfp-1* are likely responsible for cell fate specification, while *soxP-3* and *pax-2/5/8* are more likely involved in driving aspects of epidermal lineage morphogenesis.

### PROG-2-5 is a novel secreted protein that defines the Hyman vesicles, specialized organelles specific to the planarian epidermis

The molecular properties and functions of the non-rhabdite epidermal granules, generally termed rhabdoids (Hyman, 1951), are completely unknown. Their likely secretory properties make them prime candidates in modulating the chemical microenvironment at the epidermal surface of planarians. Several hypothesis have been proposed regarding the nature of these secretory vesicles. Coward and Piedelato suggested that the rhabdoids may be responsible for generating the noxious taste characteristic of Turbellarians, thus repulsing potential predators (Coward and Piedilato, 1972). Additionally, Tyler suggested that rhabdoids may contain materials that can be added to the apical surface of the epidermal cells to strengthen their barrier function, or adhesive functions (Tyler, 1976). Here we show that a member of the PROG family of small, secreted proteins (PROG-2-5) is a specific component of these vesicles. Because the term rhabdoid is currently used in the literature to describe either pediatric malignant tumors (Geller et al., 2015) or the rod-shaped protoplasmic body found in the sensitive cells of insectivorous plant leaves (Darwin, 1875), we thought it appropriate to honor the memory and meticulous work of Libbie Hyman (Hyman, 1951) by naming these PROG-2-5+ organelles Hyman vesicles.

Still, the function of these vesicles remains to be determined. Their elimination by chronic *soxP-3* dsRNA treatment (3 months) did not result in an observable defect under standard laboratory conditions, suggesting that these vesicles may have highly specialized functions associated with environmental selective pressures we have yet to recapitulate in captivity. The fact that *prog* genes exist in pairs and are all exquisitely regulated by *soxP-3* suggests that these genes have been subjected to selective pressure. Gene duplications are a molecular mechanism known to mediate the generation of functional novelty. As such, it is thought that selective pressures acting on the gene copies immediately after the duplication event shape the maintenance and functional divergence of duplicated genes (Hittinger and Carroll, 2007; Des Marais and Rausher, 2008). It still remains a distinct possibility, therefore, that *prog* genes may exist in other organisms, as they may represent fast evolving orthologs whose homology by sequence alone cannot be easily ascertained. This possibility can only be tested by first carrying out extensive taxon sampling to overcome long evolutionary distances, followed by functional studies of the identified candidates to establish orthologies. Such an approach, if systematically executed, should help further determine the extent of conservation that may or may not exist between the different strategies deployed by animals with homeostatically dynamic epidermal tissues to diversify and specialize their respective epithelial cells.

## Conclusion

Our work demonstrates that both conserved and unique transcriptional modules are integrated into a hierarchical regulatory program (Figure 7B) driving the specification and differentiation of the planarian epidermis (Figure 7C). We have shown that planarians, like mammals, require TP53, zinc-finger, and SOX proteins to modulate the progressive, postmitotic differentiation of epidermal cells. This is in contrast to other invertebrate model systems (*e.g.,* flies and nematodes) in which P53 is dispensable for adult epidermal development. We also uncovered a shared role between mammals and planarians for PAX proteins in modulating the fate of epidermal progenitors into specialized cells. In planarians, however, *pax-2/5/8* cooperates with *soxP-3* and togwtther promote the morphological differentiation of epidermal progenitors through the activation of a likely recently expanded family of novel and/or fast-evolving PROG proteins. Our study sheds light on the genetic and molecular programs that likely underpin the evolution of complex, dynamic epidermal tissues. It also puts forward the planarian epidermis as a useful invertebrate model system to study the conserved and derived mechanisms coordinating cell proliferation and differentiation in epidermal tissues undergoing constant homeostasis.

## Material and Method

### Animal care and irradiation

The CIW4 clonal line of *S. mediterranea* was maintained at 20°C in IX Montjuic salts (Cebrià and Newmark, 2005) supplemented with 50 μg/mL gentamicin sulfate and fed homogenized calf liver paste once per week. Planarians 2-3 mm in length were starved for at least 1 week before use. Irradiation was done with a GammaCell 40 Exactor irradiator at a dose rate of 85 Rad/minute.

### Gene cloning

Genes in this study were cloned from a CIW4 regeneration time-course cDNA library into the pPR-T4P vector as previously described (Gurley et al., 2008).

Primers used in this study:

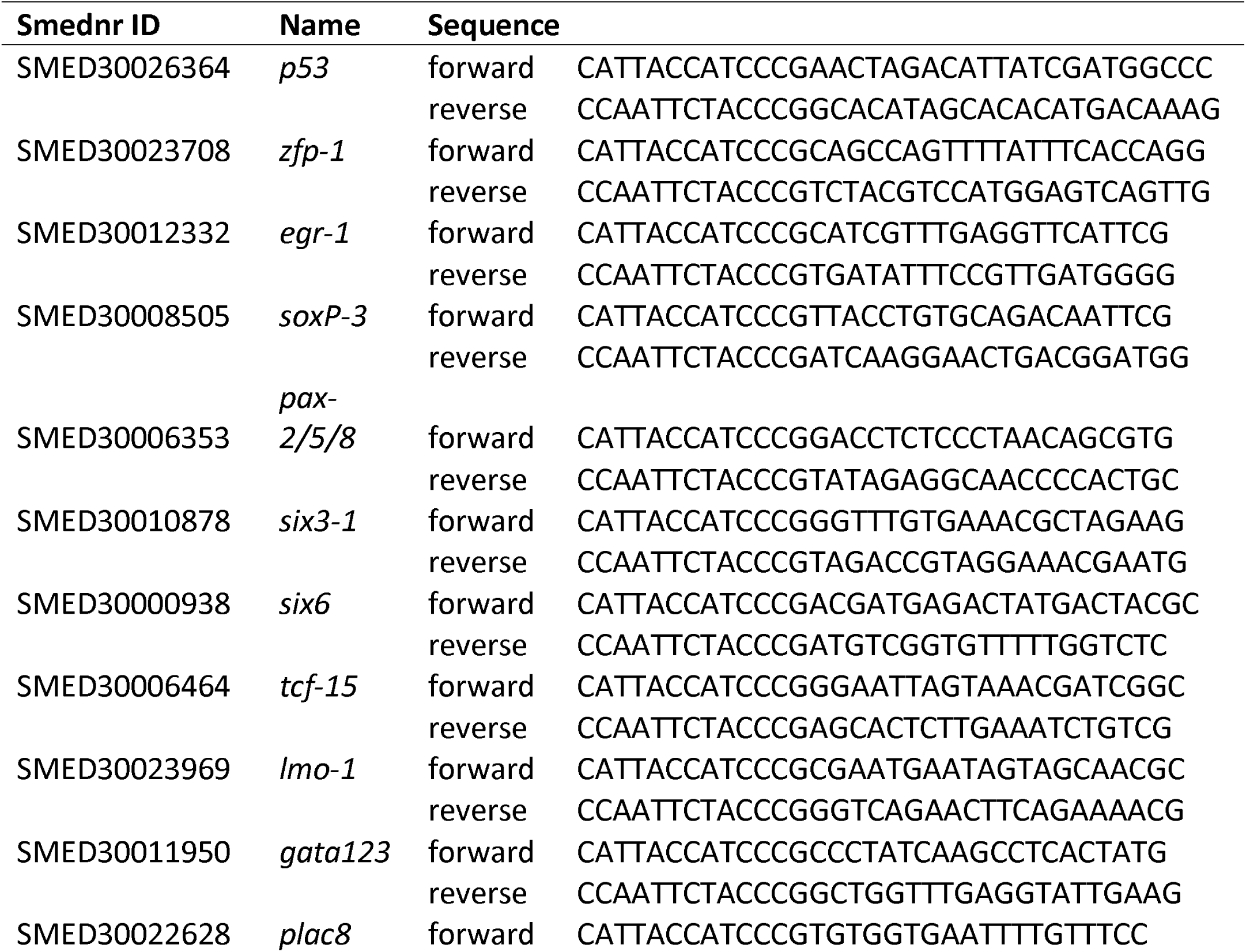
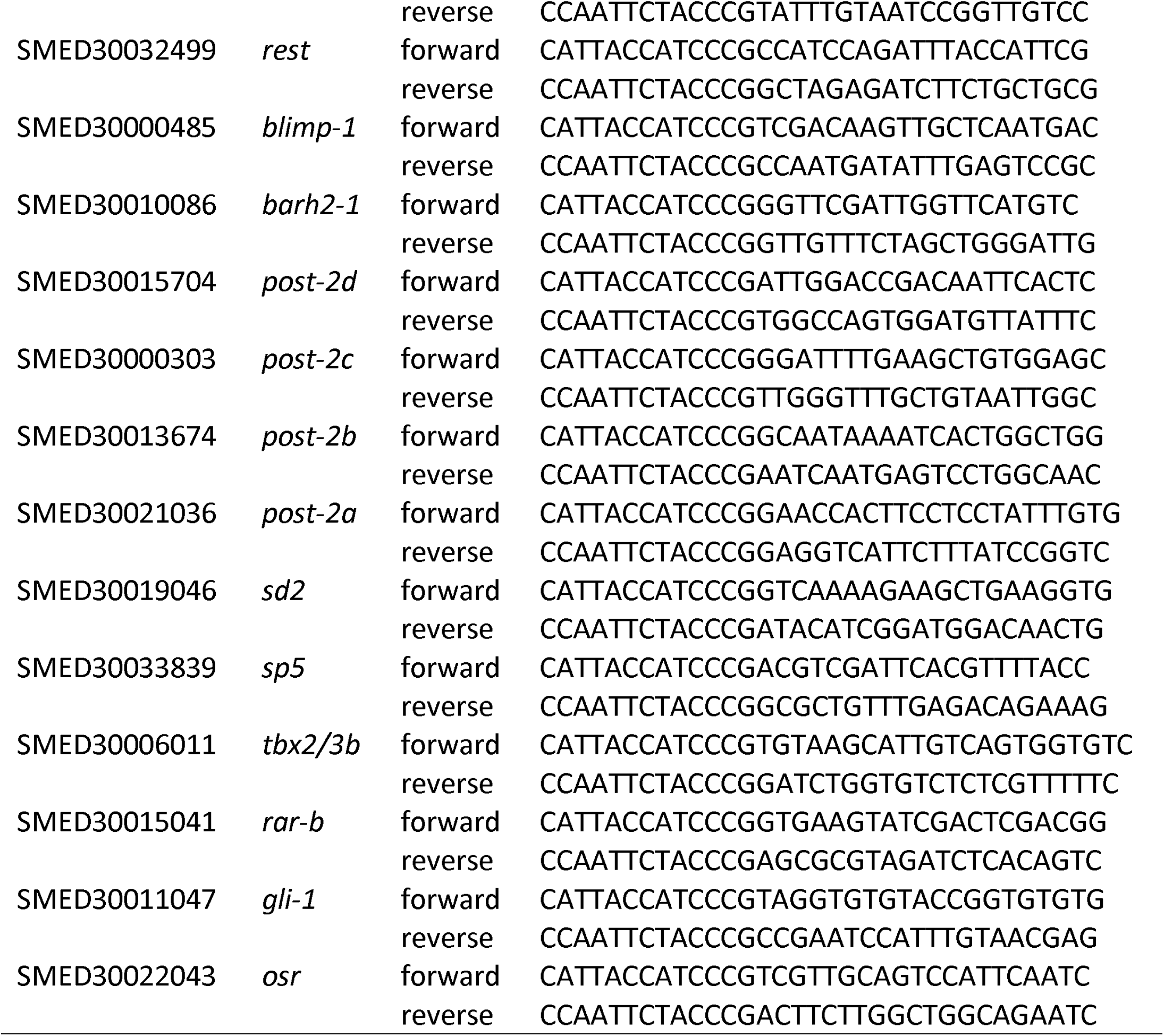

### RNAi

RNAi was performed by feeding as previously described (Gurley et al., 2008) with modifications. Briefly, cloned gene vectors were transformed into bacterial strain HT115, cultured in 2XYT to O.D.= 0.8 −1, induced to express dsRNA for 2 hours with IPTG (final 1mM) at 37°C with vigorous shaking. Bacterial pellets were rinsed once with Milli-Q water, and mixed with an equal weight of calf liver paste. dsRNA food was given to the animals every 3 days for three feedings.

### FACS

FACS sorting of X1s was performed as previously described (Reddien et al., 2005; Hayashi et al., 2006) with modifications. Briefly, animals were minced and dissociated in ice-cold CMF supplemented with 1% BSA. Cell suspension was filtered through 20 μm nylon mesh, pelleted, resuspended in room temperature IPM (Schürmann and Peter, 2001), supplemented with Hoechst 33342 (final = 50 μg/mL), and incubated at R.T. in the dark for 40 minutes. Propidium iodide was added (final = 1 μg/mL) for the last 5 minutes to label dead cells. Viable cells were then sorted using Influx (BD Biosciences).

### RNA-Seq and gene expression analysis

Whole worms or sorted cells were homogenized in TRIzol reagent (Sigma) and RNA was isolated as described in the manufacturer’s manual. RNA from whole worms was treated with RNase-free DNase on QIAGEN RNAeasy columns and eluted in nuclease-free water. For each replicate, 1 μg RNA from five worms or 100 ng RNA from 10^5^ sorted cells was used to generate RNA-Seq libraries using the Illumina TruSeq Stranded kit. Libraries were sequenced in 50bp single reads (whole worms) or in 150bp paired reads (sorted cells) using the Illumina HiSeq 2500 sequencer. RNA-Seq analysis was carried out by mapping reads to the transcriptome as previously described (version: smed_20140614) (Robb et al., 2015; Tu et al., 2015). Reads were aligned using Bowtie with the following parameters: -best -strata -v 2 -m 5, and read counts to genes were tallied from the SAM files with a custom script. Differential gene expression was evaluated using R and the edgeR library (Robinson et al., 2010). P-values were adjusted, as previously described (Benjamini and Hochberg, 2000). Analyses were based on the mean expression values of four biological replicates, except for the *p53(RNAi)* whole worm dataset in which 3 replicates were used (Tu et al., 2015). Fold changes were normalized and compared to controls of the same time-point (*unc-22* dsRNA treated animals). The RNA-Seq data and the transcriptome against which it was quantified are archived at GEO (Edgar et al., 2002) and are available under accession number: GSE80562.

### GO-enrichment analysis

*S. mediterranea* transcripts were aligned with NCBI BLASTX to Swiss-Prot protein database (Consortium, 2015). The top BLAST hits with a minimum e-value of 0.001 were used to annotate transcripts. GO functional categories were assigned and the enrichment was generated using DAVID (version 6.7) (Huang et al., 2009). Terms were clustered using ReVigo script (Supek et al., 2011) at a similarity cutoff of 0.7 (Medium). A representative term from each cluster was plotted based on GO term enrichment p-value.

### Transcription factors homology predictions

Possible transcription factors were identified by domain annotation as well as manual curation. PFAM domains were identified in target sequences using HMMER search (Eddy, 1998; Finn et al., 2016) with an e-value cutoff of 0.01. The list was then compared to another list created from those annotated by the Gene Ontology Consortium with GO term “regulation of transcription” (GO: 0006355) (Ashburner et al., 2000).

### *In situ* hybridization, immunostaining, and TUNEL assay

Anti-sense riboprobes were made by *in vitro* transcription with T7 polymerase from genes in the pPR-T4P vector as previously described (Pearson et al., 2009). Whole mount *in situ* hybridization (WISH) and fluorescent *in situ* hybridization (FISH) were performed as previously described (King and Newmark, 2013).

Immunostaining of anti-H3P (dilution 1:2000, Millipore, 63-1C-8) was performed after whole-mount FISH, and was detected with Alexa Fluor-conjugated antibodies (dilution 1:1000, Invitrogen). Both antibody incubations were performed for overnight at 4°C.

Immunostaining of anti-PROG-2-5 was performed on cryosections following FISH. Briefly, 4-6 mm planarians were rinsed with ice-cold 1X Montjuic salt, fixed in ice-cold 4% formaldehyde/PBSTx (0.3% Triton-X) for overnight, rinsed with PBSTx, infiltrated with 30% sucrose/PBS, embedded in O.C.T, frozen on dry ice and cut into 10 μm sections. After cryo-sectioning, samples were rinsed in PBS, bleached with 3% H_2_O_2_/PBS for 30 minutes under light, rinsed with PBS, equilibrated with 50% hybridization buffer, and hybridized with appropriate riboprobes (0.1-0.2 μg/mL) at 60°C for overnight on a humidified horizontal tray. Excess riboprobes were then removed by stringent washes in the 15-mL slide mailer (1X with 50% formamide/5X SSC, 3X with 2X SSC, 3X with 0.2X SSC, all done in 60°C with 20 minutes intervals). Samples were rinsed with PBS, blocked with 5% hose serum (Sigma) and 0.5% Western blocking reagent (Roche) in PBSTw (0.1% Tween-20), incubated with appropriate HRP-conjugated antibodies diluted in PBSTw at R.T. for 2 hours (antibody dilutions: anti-DIG, 1:500; anti-DNP, 1:200, anti-FITC, 1:500). Antibodies were washed with PBSTw and developed with tyramide-conjugated fluorophores diluted in 0.006% H202/borate buffer. After desired signals were developed, HRP activity was quenched with 200mM sodium azide. Immunostaining of anti-PROG-2-5 was performed at 4°C for overnight (1:5000 diluted in PBSTx), washed, and incubated with Alexa Fluor-conjugated secondary antibodies (1:1000 diluted in PBST) for 2 hours at R.T.

TUNEL assay was performed as previously described (Pellettieri et al., 2010; Tu et al., 2015).

### Antibody production

Full length *prog-2-5* was cloned into pET21a vector for recombinant protein induction in *E. coli.* Rabbit polyclonal antibodies were generated against purified recombinant PROG-2-5 and affinity purified (Proteintech).

### Western blotting

Planarian total lysate was extracted in ice-cold 2X lysis buffer (4% SDS, 200mM DTT, 250mM Tris-HCl pH6.8, supplemented with 2X cOmplete ULTRA (Roche) prior use) and centrifuged at full speed (>10,000 g) for 15 minutes at 4°C. The supernatant was mixed with an equal volume of 2X sample buffer, boiled for 5 minutes, loaded on 4-20% Mini-Protean TGX gels (BioRad), and transferred to 0.45 μm PVDF membranes. The membranes were blocked in 5% non-fat milk/TBSTw (0.1% Tween-20), incubated with appropriate primary antibodies (diluted in blocking reagent) overnight at 4°C with shaking, washed with TBSTw, incubated with appropriate HRP-conjugated secondary antibodies (diluted in TBSTw) for 1 hour at R.T., washed and developed using Amersham ECL Western Blotting Detection Reagent or SuperSignal West Dura Extended Duration Substrate (ThermoFisher). Antibody dilutions as follow: Rabbit anti-PROG-2-5 polyclonal IgG 1:10,000; Mouse anti-α Tubulin monoclonal IgG (Sigma, T5168) 1:10,000.

### Electron microscopy

For ultrastructural analysis, worms were fixed in cold fixative containing 2.5% glutaraldehyde and 2% paraformaldehyde in cacodylate buffer (pH 7.35) and stored at 4°C in fixative until processing. After rinse in the same buffer, samples were fixed in 1% Osmium tetroxide 2 hours and En Bloc staining overnight in 0.5% aqueous uranyl acetate. Samples were then dehydrated in gradient acetone at 4°C, infiltrated at room temperature with Spurr’s resin (Electron Microscopy Sciences Kit and recipe), and polymerized at 60°C for 48 hours. Thin sections were cut at 50 nm on a Leica UC6 ultramicrotome,collected on slot grids, and post-stained in Sato’s lead and 4% uranyl acetate.

For immunogold labeling, planarians were fixed in 4% paraformaldehyde and 0.1% glutaraldehyde in 0.1 M PBS (pH 7.4) for overnight, dehydrated in gradient ethanol, and infiltrated and embedded in LR White resin. Sections of about 70nm were stained with rabbit anti-PROG-2-5 primary antibody (1:200) followed by 12 nm colloidal gold goat-anti-rabbit secondary antibody (Jackson Immunoresearch Labs; 1:25 dilution). All images were taken on a FEI Technai BioTwin microscope equipped with a Gatan Ultrascan 1000 digital camera.

### *De novo* motif analysis

The *S. mediterranea* transcriptome (version: smed_20140614) were aligned to the *S. mediterranea* sexual genome assembly version 3.1 (SmedSxl_v3.1) using map2assembly, a script included in the MAKER package (v2.31.8) (Holt and Yandell, 2011). The alignment for each gene of interest with the best map2assembly score was used to pinpoint the genomic sequence regions for which to extract promoters, 2kb or 5kb upstream of the coding sequence.

To identify putative regulatory regions, the MEME Suite (4.11.1) (Bailey et al., 2015) was used. Two sets of upstream sequences (2kb), the experimental set (13 sequences from genes down-regulated in both *soxP-3-* and *pax-2/5/8(*RNAi) animals) and the background set (32,104 sequences from all *S. mediterranea* genes mapped) were first processed with psp-gen to generate a position-specific prior that was then used as input to MEME. TOMTOM was run with the JASPER motif database to identify consensus to known motifs (Gupta et al., 2007; Mathelier et al., 2016).

To determine the p63 motifs, p63scan program was downloaded from http://131.174.198.125/bioinfo/p53scan/ and supplied a FASTA file with the promoter sequences (5kb) for 19 TFs (of which expression levels are down-regulated in both *p53-* and *zfp-1* (RNAi) X1s) using optimal threshold as described previously (Kouwenhoven et al., 2010).

## Acknowledgements

We would like to thank members of the Sánchez Alvarado lab for critical reading of the manuscript, especially Sarah Elliot for the editing of our first draft. We thank Hanh Vu and Erin Davies for kindly sharing their plasmids and RNA probes. We thank Eric Ross and Kirsten Gotting for bioinformatics assistance, Sean McKinney for image processing assistance, Melania McClain and Kexi Yi for EM assistance, Allison Peak for RNA-seq library assistance, Mark Miller for illustration assistance, and all members of the Cytometry, Histology, Molecular Biology and Planaria core facilities at the Stowers Institute for their technical support. This work was supported by NIH R37GM057260 to ASA. ASA is an investigator of the Howard Hughes Medical Institute and the Stowers Institute for Medical Research.

## Additional Information

Original data underlying this manuscript can be accessed from the Stowers Original Data Repository at http://www.stowers.org/research/publications/LIBPB-1174

## Author contributions

LCC, ASA, conception and design, drafting and revising the article; LCC, KTU, CWS, SMR, FLG, acquisition of data, and analysis; LCC, KTU, CWS, SMR, FLG, ASA interpretation of data.

## Supplementary figures

**Figure 1 Supplement 1.**
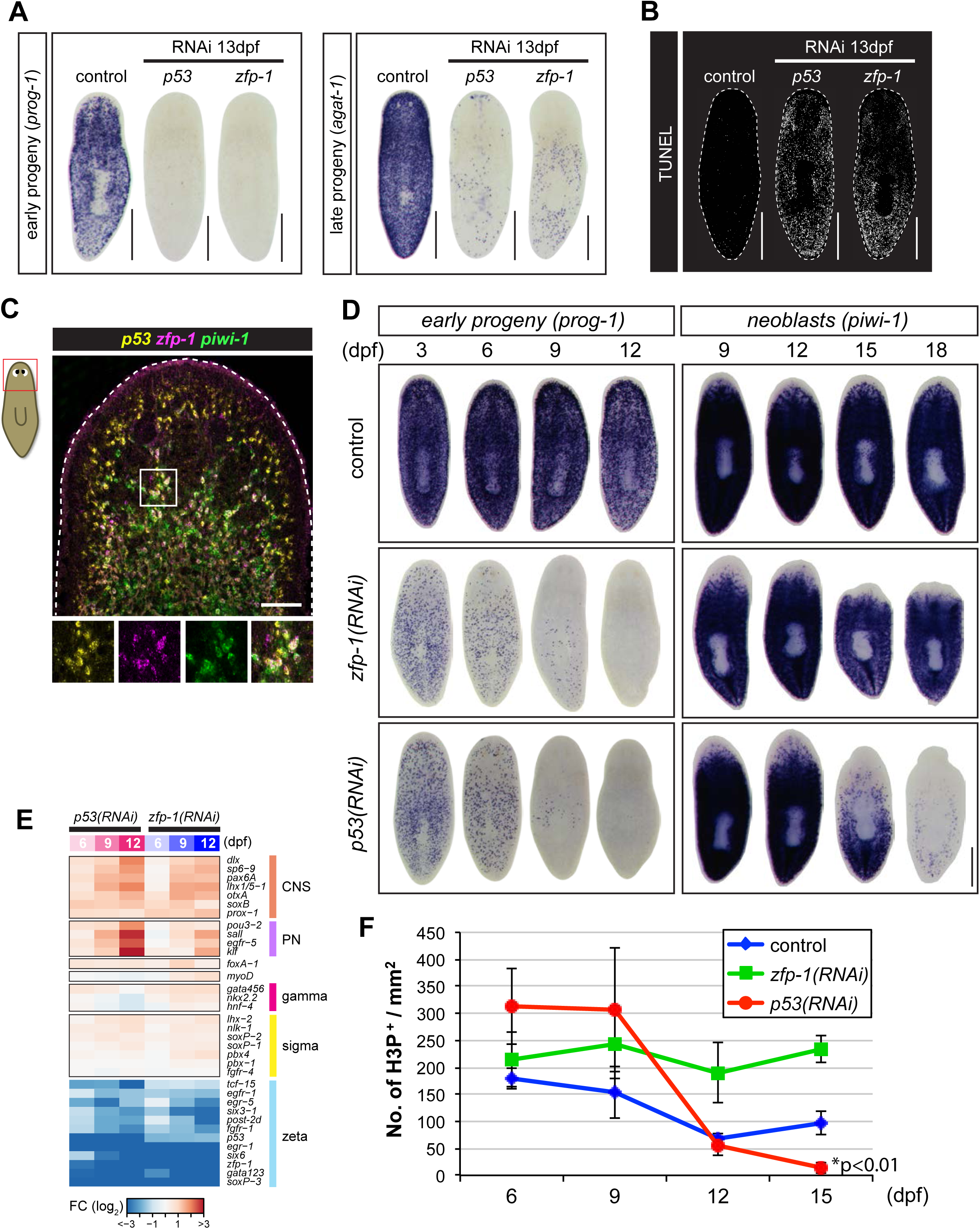
Progression of *p53-* and *zfp-1 (RNAi)* phenotypes. (A) *p53* and *zfp-1* RNAi cause complete loss of early progeny *(prog-1*)* and late progeny *(agat-1*)* cells monitored by whole-mount *in situ* hybridization (WISH). Dpf, days post dsRNA feeding. Scale bar, 500μm. (B) *p53* and *zfp-1* RNAi induce massive apoptosis assessed by whole-mount TUNEL assay (n >8 per RNAi, 100% penetrance). Scale bar, 500μm. (C) Fluorescent *in situ* hybridization (FISH) of *p53, zfp-1,* and *piwi-1* riboprobes detects co-localization of *p53* and *zfp-1* in the neoblast compartment (*piwi-l*^*+*^). Boxed region is split into individual channel shown at the bottom. Scale bar, 100μm. (D) Defects in epidermal differentiation (loss of *prog-1*^*+*^ cells) are detected in both *p53-* and *zfp-1 (RNAi)* animals as early as 3 dpf. Defects in neoblast maintenance (loss of *piwi-l*^*+*^ cells) is observed only in *p53(RNAi)* animals after 15 dpf and not in *zfp-1 (RNAi)* animals. Scale bar, 500μm. (E) Lineage markers of central nervous system (CNS) and protonephridial system (PN) are up-regulated in *p53-* and *zfp-1 (RNAi)* X1 cells, whereas genes expressed in zeta-class neoblasts (epidermal lineage) are down-regulated. Lineage markers of pharynx *(foxA*), muscle *(myoD),* gut (gamma) and sigma-class neoblasts are not significantly changed. (F) Immunostaining of anti-phospho-Histone 3 (H3P) shows decline (*p<0.01) of cell proliferation in *p53(RNAi)* animals (mean ± s.d., n > 8 per time point).

**Figure 1 Supplement 2.**
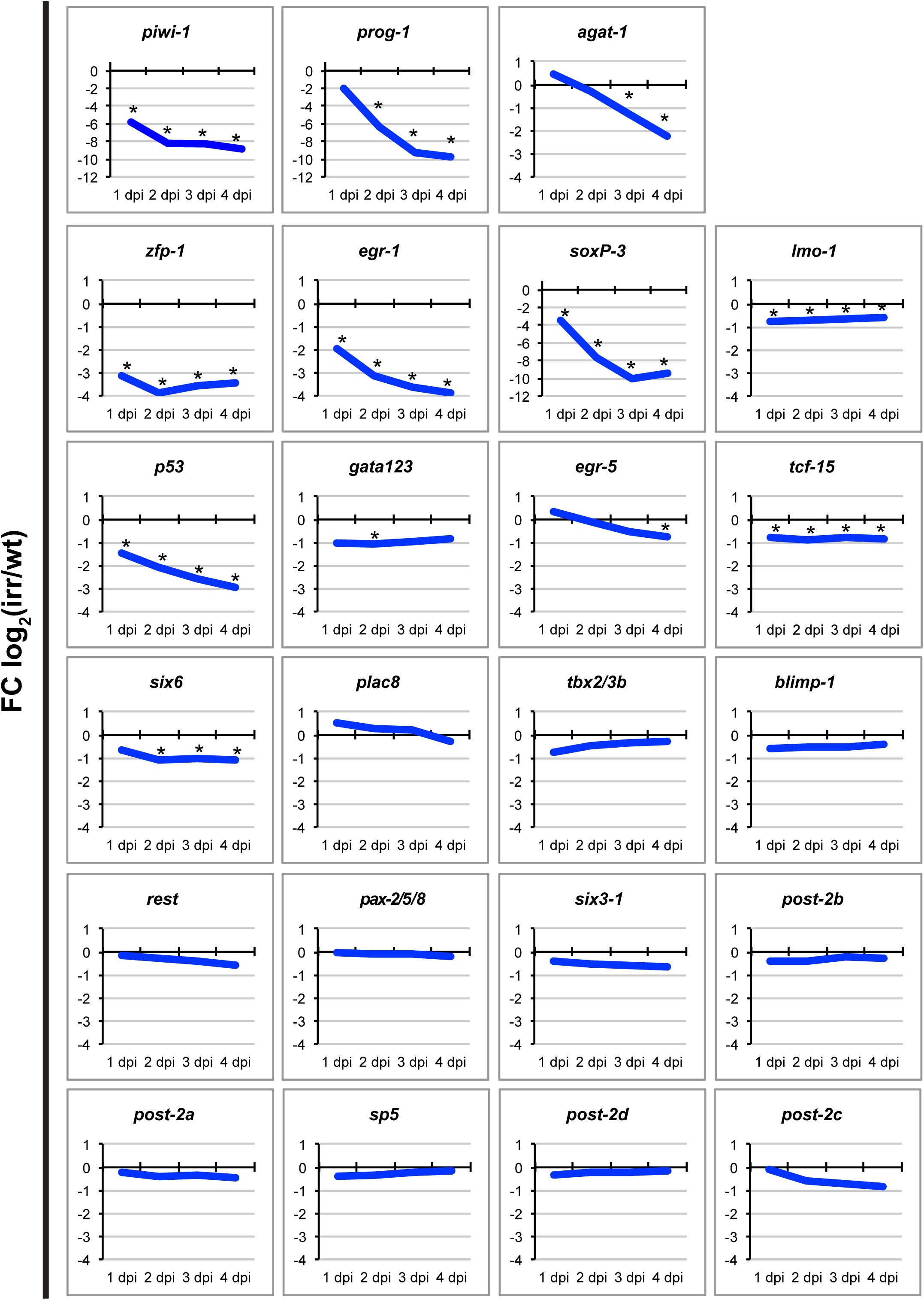
Expression levels of candidate TFs in irradiated animals. Decreased expression upon irradiation is observed for several TFs downstream of *p53* and *zfp-1* analyzed by RNA-Seq. Notably, not all TFs downstream of *p53* and *zfp-1* show significant down-regulation in the irradiated animals, likely due to their higher expression in the mature epidermis or in other tissue. Dpi, days post irradiation. Asterisks, p-adj < 0.01.

**Figure 2 Supplement 1. Expression of late progeny markers are generally not affected by *soxP-3* RNAi** WISH of late progeny markers *pmp-6, pmp-10, X1.C3.3, zpuf-2,* and *zpuf-*3. Scale bar, 500μm.

**Figure 3 Supplement 1.**
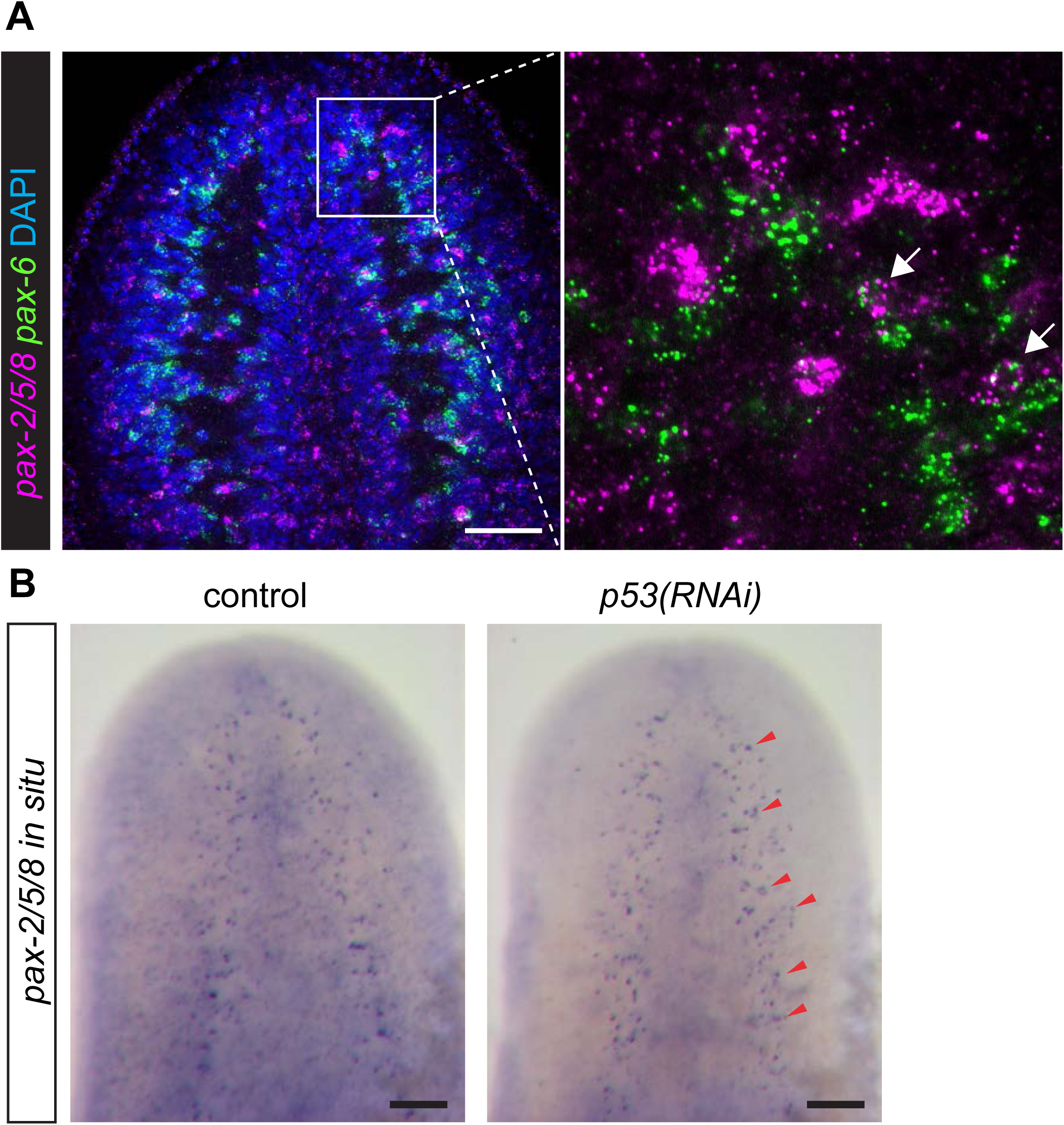
*pax-2/5/8* expression in the CNS is independent of *p53* function. (A) *pax-2/5/8* and *pax-6* double FISH in the cephalic ganglia. Higher magnification of the boxed region is shown in the right panel. Arrows point to *pax-2/5/8*^*+*^*/pax-6*^*+*^ double positive cells. Scale bar, 100μm. (B) *pax-2/5/8* WISH signals in the cephalic ganglia is not affected by *p53* RNAi (arrowheads). Scale bar, 100μm.

**Figure 3 Supplement 2.**
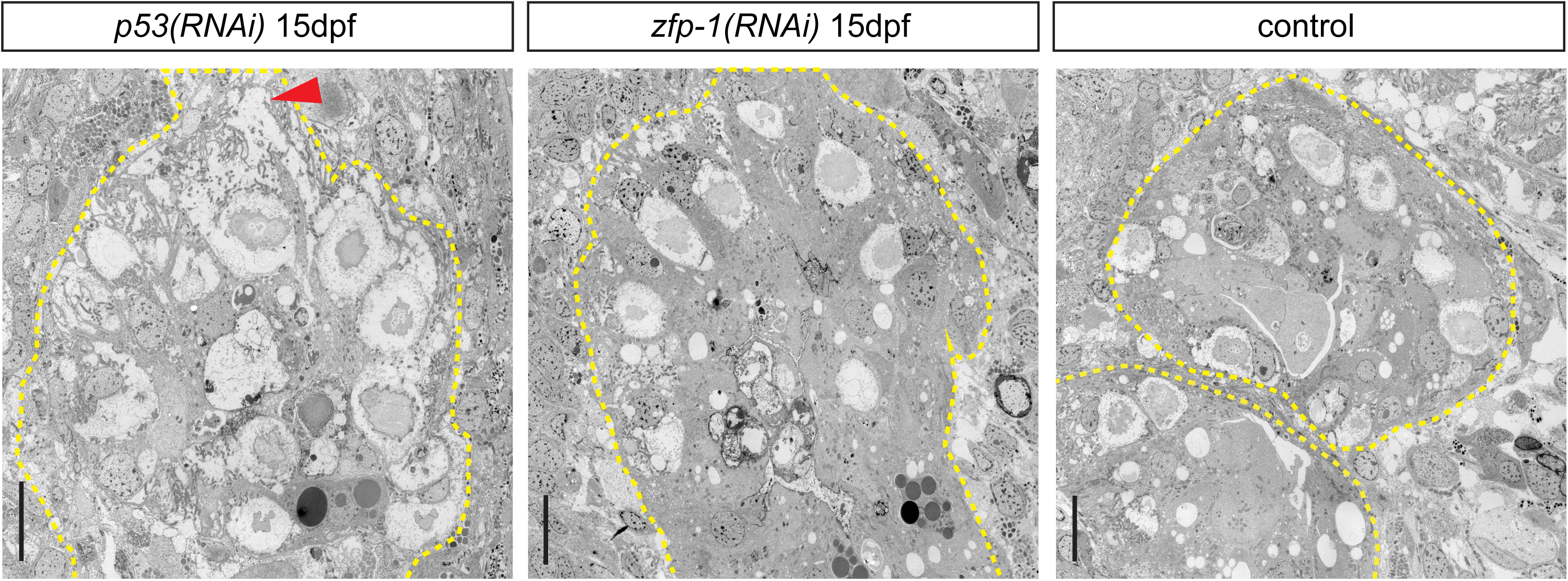
*p53(RNAi)* animals show morphological defects in the gut. TEM micrographs of gut sectioned at the tertiary branch (outlined by dash lines). *p53(RNAi)* gut cells show severe morphological defects (arrowhead point to a rupture), n = 2,100% penetrance. Scale bar, 10μm.

**Figure 4 Supplement 1.**
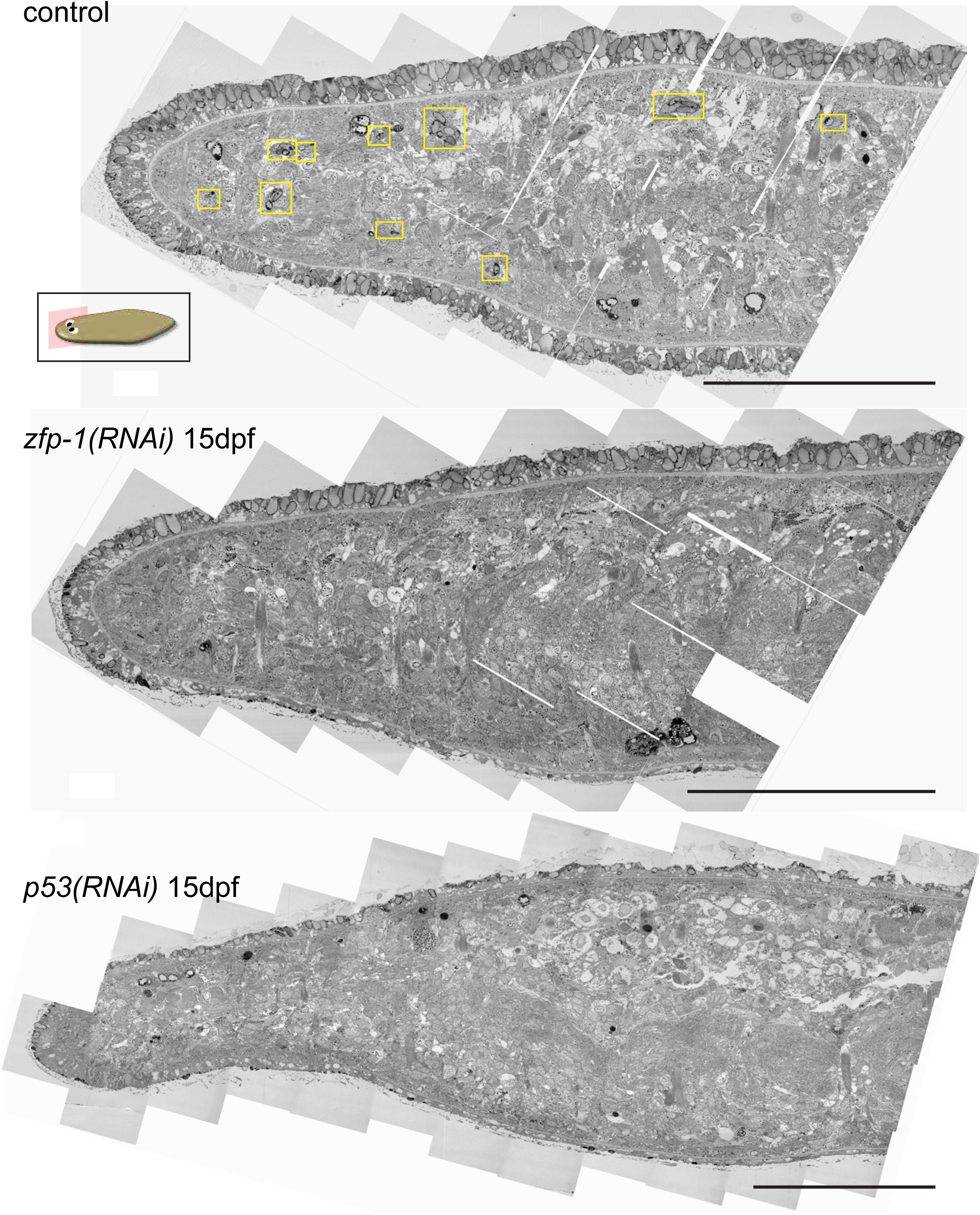
*p53-* and *zfp-1* RNAi reduce the number of rhabdite-forming cells in the mesenchyme. Tiled TEM micrographs of entire head region. Rhabdite-forming cells (boxes in control) are eliminated in *p53-* and *zfp-1 (RNAi)* animals, n = 3, 100% penetrance. Scale bar, 100μm.

**Figure 5 Supplement 1.**
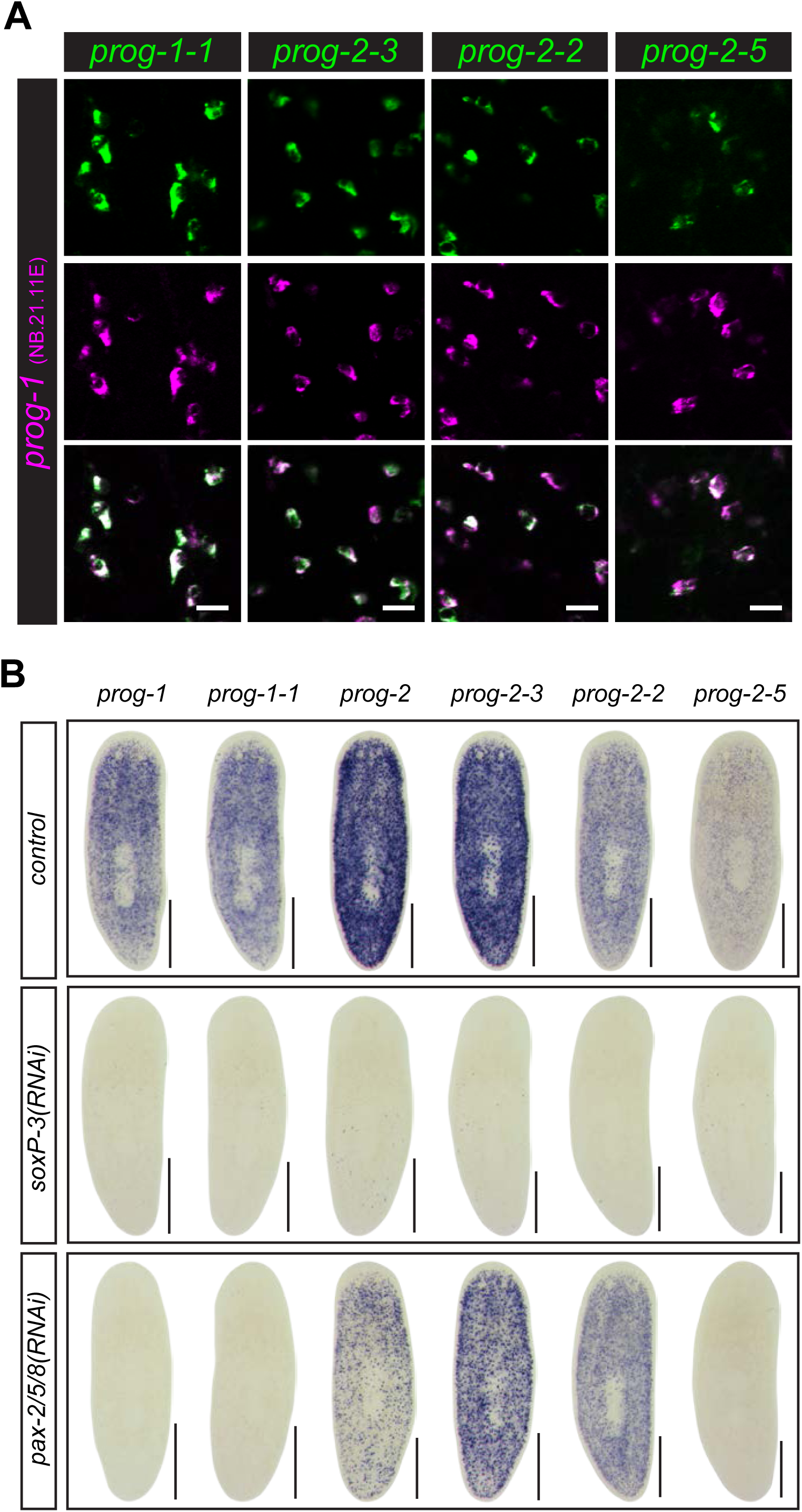
Expression patterns of *prog* gene family members. (A) Double FISH of *prog-1* with *prog-1-1, prog-2-3, prog-2-2* and *prog-2-5* show near 100% of co-localization. Scale bar, 20μm. (B) Expression of *prog* genes in *soxP-3-* and *pax-2/5/8(RNAi)* animals analyzed by WISH. Note that *prog-2-2* is not significantly down-regulated in *pax-2/5/8* RNAi. Scale bar, 100μm.

**Figure 5 Supplement 2.**
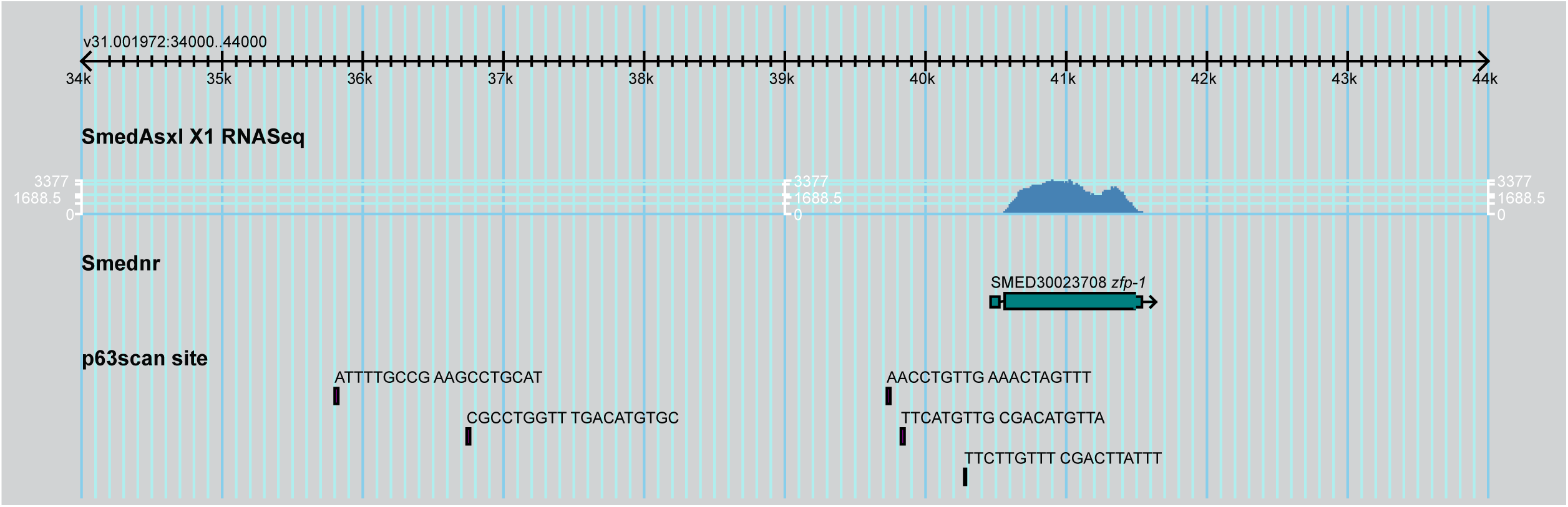
Vertebrate TP63 binding sites are predicted in the upstream sequence of *zfp-1* locus in *S. mediterranea* genome. Screenshot of SmedGD browser showing 5kb upstream sequence of *zfp-1* locus in *S. mediterranea* genome V3.1. Five TP63 binding sites are predicted by p63scan (Kouwenhoven et al., 2010).

**Figure 6 Ssupplement 1.**
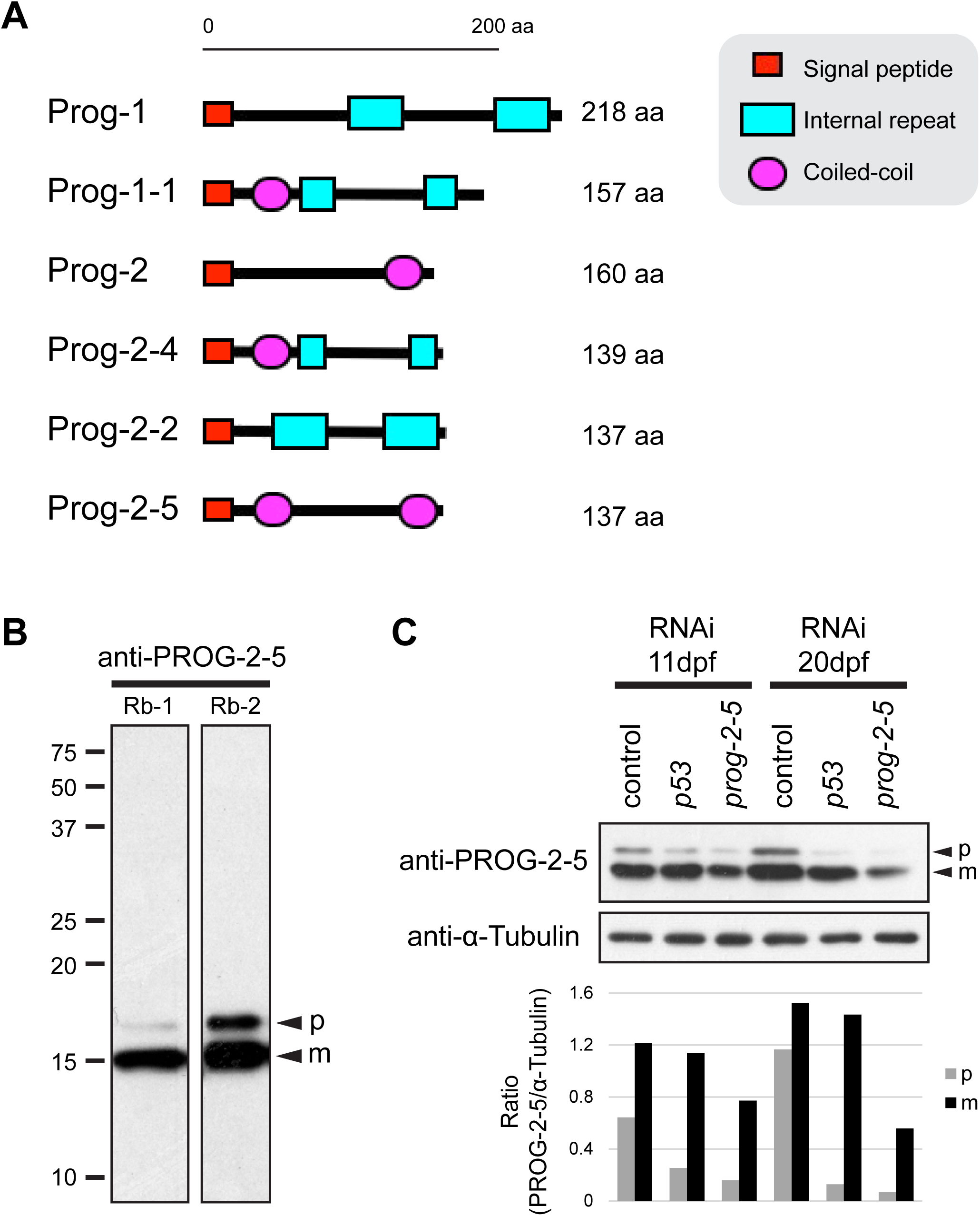
Characterization of anti-PROG-2-5 antibodies. (A) SMART analysis (Schultz et al., 1998) of Prog gene family predicts signal peptide, internal repeats, and coiled-coil region in their protein sequences. (B) Western blot analysis of PROG-2-5 proteins. Two polyclonal antibodies (Rabbit-1 and Rabbit-2) raised against full-length PROG-2-5 protein specifically detect two bands corresponding to the precursor (p) and mature (m) forms of PROG-2-5 proteins in the planarian whole worm lysates. (C) Western blot analysis of PROG-2-5 in *p53-* and *prog-2-5(RNAi)* whole worm lysates. Larger bands corresponding to the precursor form of PROG-2-5 proteins (p) are knocked-down in both *p53-* and *prog-2-5(RNAi)* lysates. However, mature form of PROG-2-5 proteins (m) are only significantly decreased in *prog-2-5(RNAi).* Relative densities of PROG-2-5 bands (Rabbit-2) to α-Tubulin are shown at the bottom.

**Figure 6 Supplement 2.**
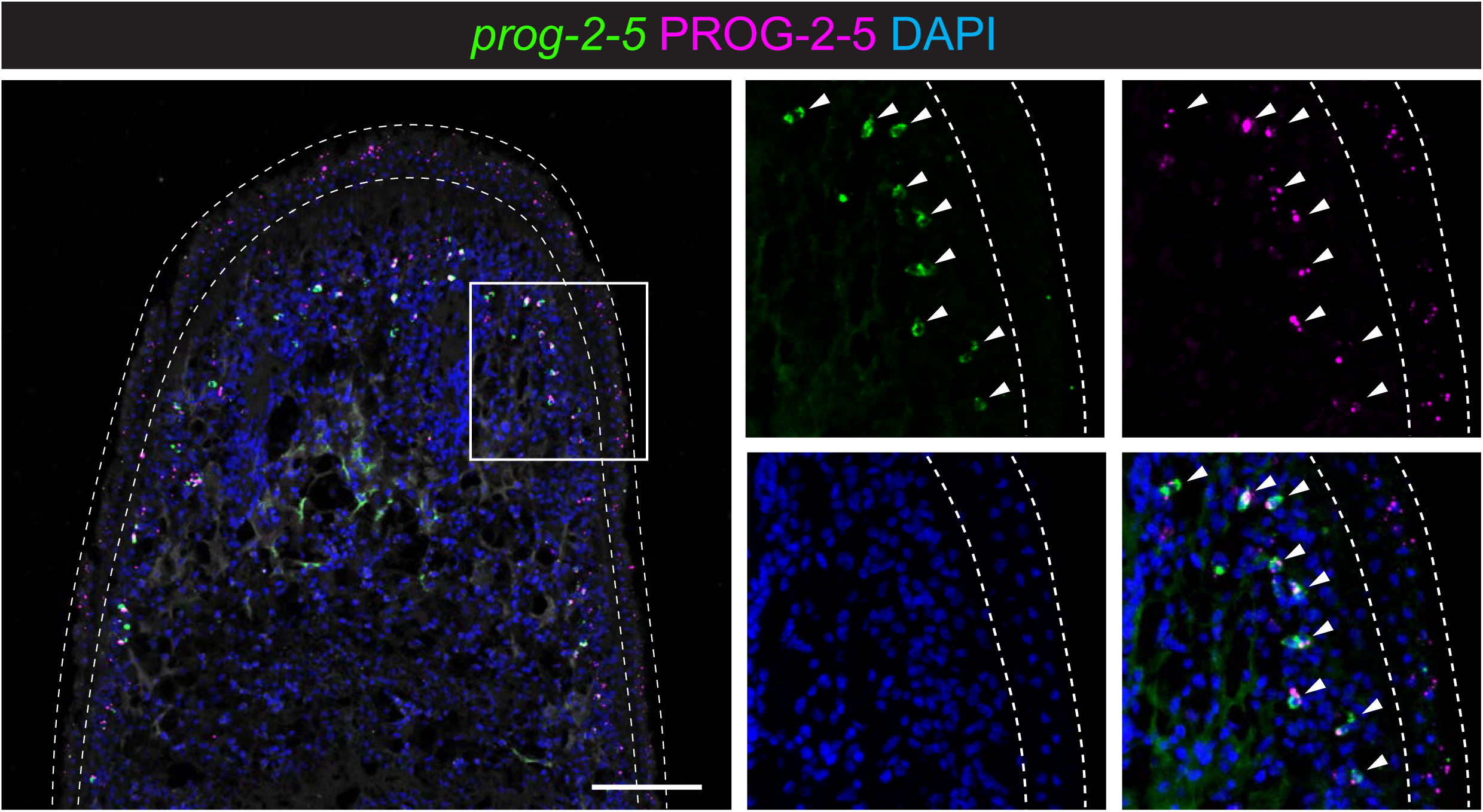
PROG-2-5 proteins localize to the mature epidermis. Immunostaining of PROG-2-5 (Rabbit-2) combining FISH of *prog-2-5* detects bright PROG-2-5^+^ puncta in early progeny cells (*prog-2-5*^*+*^) as well as in the epidermis (*prog-2-5*^*ne*^*g*, outlined by dash lines). Higher magnification of the boxed region is split into individual channels in the right panels. Scale bar, 100μm.

**Figure 6 Supplement 3.**
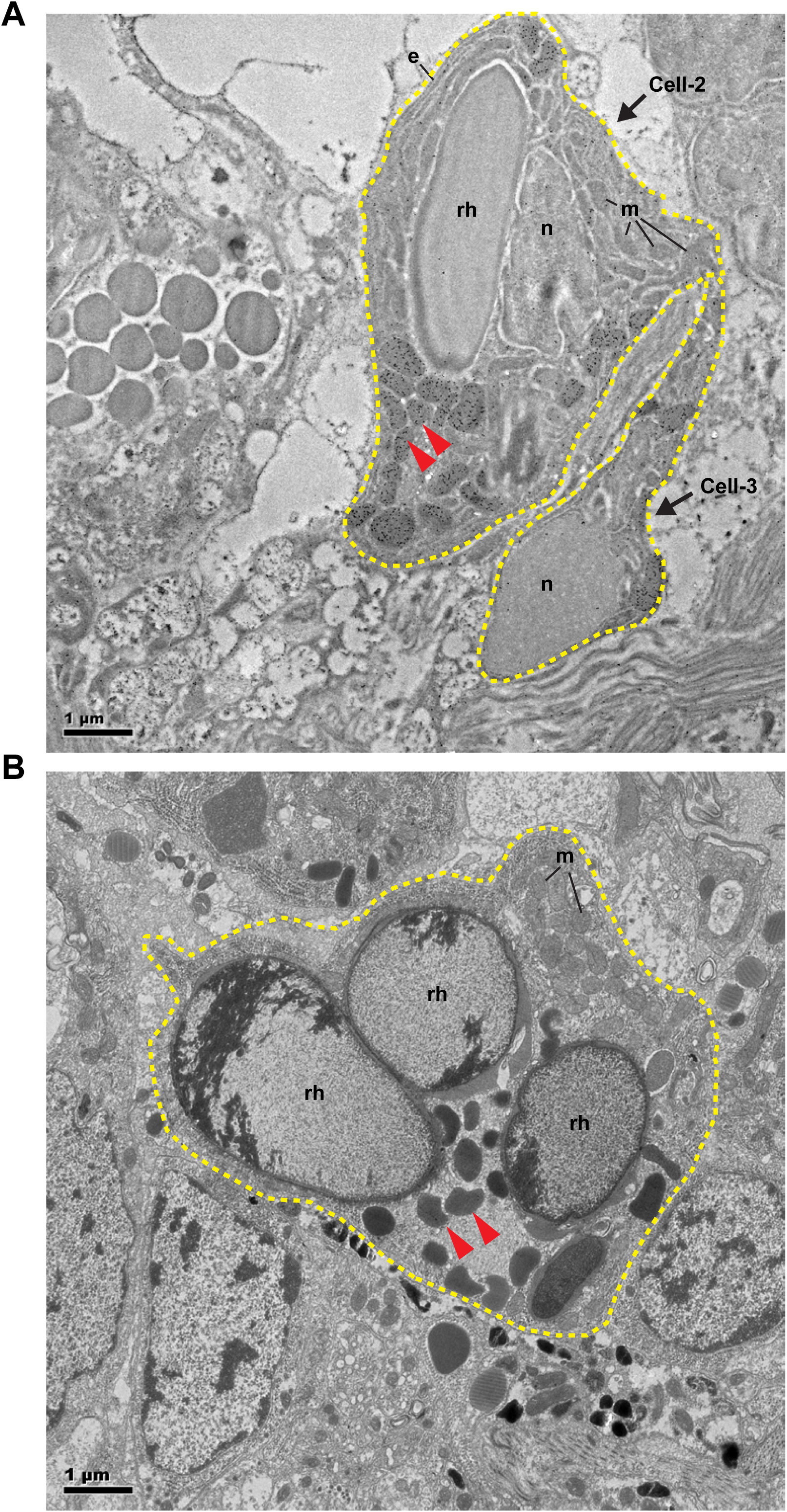
PROG-2-5 proteins are detected in the oval-shape granules of rhabdite-forming cell. (A) IEM image of PROG-2-5^+^ mesenchymal cells (outlined by dash lines). Arrowheads point to the PROG-2-5^+^ granules, e, endoplasmic reticulum, n, nucleus, m, mitochondria, rh, rhabdite. Scale bar, 1μ.m. (B) TEM image with the same magnification of (A) showing a rhabdite-forming cell (outlined by dash line). Arrowheads point to the electron-dense granules. Scale bar, lμm.

## Supplementary files

Supplementary Table 1. Summary of 596 genes significantly changed in *p53-* or *zfp-1 (RNAi)* X1 cells

Supplementary Table 2. GO term enrichment of 208 common DE-genes in *p53-* and *zfp-1 (RNAi)* X1 cells

Supplementary Table. Expression profiles of candidate TFs in the irradiation time-course analyzed by RNA-Seq

Supplementary Table 4. Summary of expression patterns of candidate TFs analyzed by FISH

Supplementary Table 5. Summary of 594 DE-genes identified in the *pax-2/5/8-, soxP-3-, zfp-1-* or *p53(RNAi)* animals

Supplementary Table 6. Genome coordinates and expression levels (RPKM) of *soxP-3*- and *pax-2/5/8(RNAi)* common DE-genes

Supplementary Table 7. Genome coordinates of candidate TFs and the number of TP63 binding sites predicted by p63scan

